# Epistasis at the SARS-CoV-2 RBD Interface and the Propitiously Boring Implications for Vaccine Escape

**DOI:** 10.1101/2021.08.30.458225

**Authors:** Nash D. Rochman, Guilhem Faure, Yuri I. Wolf, Peter L. Freddolino, Feng Zhang, Eugene V. Koonin

## Abstract

At the time of this writing, December 2021, potential emergence of vaccine escape variants of severe acute respiratory syndrome coronavirus 2 (SARS-CoV-2) is a grave global concern. The interface between the receptor-binding domain (RBD) of SARS-CoV-2 spike (S) protein and the host receptor (ACE2) overlap with the binding site of principal neutralizing antibodies (NAb), limiting the repertoire of viable mutations. Nonetheless, variants with multiple mutations in the RBD have rose to dominance. Non-additive, epistatic relationships among RBD mutations are apparent, and assessing the impact of such epistasis on the mutational landscape is crucial. Epistasis can substantially increase the risk of vaccine escape and cannot be completely characterized through the study of the wild type (WT) alone. We employed protein structure modeling using Rosetta to compare the effects of all single mutants at the RBD-NAb and RBD-ACE2 interfaces for the WT, Delta, Gamma, and Omicron variants. Overall, epistasis at the RBD interface appears to be limited and the effects of most multiple mutations are additive. Epistasis at the Delta variant interface weakly stabilizes NAb interaction relative to ACE2 interaction, whereas in the Gamma variant, epistasis more substantially destabilizes NAb interaction. Although a small, systematic trend towards NAb destabilization not observed for Delta or Gamma was detected for Omicron, and despite bearing significantly more RBD mutations, the epistatic landscape of the Omicron variant closely resembles that of Gamma. These results suggest that, although Omicron poses new risks not observed with Delta, structural constraints on the RBD hamper continued evolution towards more complete vaccine escape. The modest ensemble of mutations relative to the WT that are currently known to reduce vaccine efficacy is likely to comprise the majority of all possible escape mutations for future variants, predicting continued efficacy of the existing vaccines.

**Significance:** Emergence of vaccine escape variants of SARS-CoV-2 is arguably the most pressing problem during the COVID-19 pandemic as vaccines are distributed worldwide. We employed a computational approach to assess the risk of antibody escape resulting from mutations in the receptor-binding domain of the spike protein of the wild type SARS-CoV-2 virus as well as the Delta, Gamma, and Omicron variants. At the time of writing, December, 2021, Omicron is poised to replace Delta as the dominant variant worldwide. The efficacy of the existing vaccines against Omicron could be substantially reduced relative to the WT and the potential for vaccine escape is of grave concern. Our results suggest that although Omicron poses new evolutionary risks not observed for the Delta variant, structural constraints on the RBD make continued evolution towards more complete vaccine escape unlikely. The modest set of escape-enhancing mutations already identified for the wild type likely include the majority of all possible mutations with this effect.

## Introduction

When severe acute respiratory syndrome coronavirus 2 (SARS-CoV-2), first emerged as a global public health concern early in 2020, there was considerable debate regarding whether the low mutation rate of the virus and the relatively inflexible receptor-binding domain (RBD) of the antigenic spike (S) protein would admit robust host adaptation(1, 2). By 2021, it became clear that SARS-CoV-2 has access to a broad mutational repertoire enabling extensive diversification(3) and that without vaccination, SARS-CoV-2 would likely result in substantial global disease burden for a protracted period(4, 5). The development of multiple, effective vaccines against SARS-CoV-2(6) make it possible to dramatically reduce this burden. However, at the time of writing, December, 2021, the majority of the global population remains unvaccinated as the Omicron variant is poised to replace the Delta variant as the dominant strain worldwide. Existing vaccine efficacy against the Omicron variant might be substantially reduced relative to the WT https://www.cdc.gov/coronavirus/2019-ncov/science/science-briefs/scientific-brief-omicron-variant.html, and the potential for continued evolution towards more complete vaccine escape(7) is a major global concern https://www.cdc.gov/coronavirus/2019-ncov/variants/variant-info.html.

The interface between the receptor-binding domain (RBD) of the S protein and the host receptor (ACE2) largely overlaps with the binding sites for the most potent neutralizing antibodies (NAb)(8, 9), limiting the scope of viable mutations. Nevertheless, multiple variants containing single mutations in the RBD that, to different extents, reduce NAb binding have begun to circulate(8-10). Moreover, variants with multiple mutations in the RBD have risen to dominance outcompeting the wild type (WT, identical to Wuhan-Hu-1) and single mutants (see below). These dynamics could result from non-additive, epistatic interactions among the mutated sites(10, 11) or simply from additive effects of multiple mutations(11). The effects of all single mutations in the RBD relative to the WT have been studied, and several mutations producing partial antibody escape have been identified (8, 12). Epistasis among RBD mutations has the potential to substantially increase the risk of escape variant emergence and cannot be characterized through the study of the WT alone.

Using the Rosetta software suite https://rosettacommons.org(13), we estimated and compared the effects of all single non-synonymous mutants at the RBD-NAb and RBD-ACE2 interfaces for the WT as well as Delta (452R, 478K), Gamma (417T, 484K, 501Y), and Omicron (339D, 371L, 373P, 375F, 417N, 440K, 446S, 477N, 478K, 484A, 493R, 496S, 498R, 501Y, 505H) variants. The Delta and Gamma variants were dominant in different regions of the world with rising frequencies as of Summer, 2021(14, 15). Delta rose to global dominance in the following months and, at the time of writing, Omicron is rapidly growing in frequency and is expected to become the next globally dominant strain. We establish the distribution of RBD mutations on the plane bounded by the costs of ACE2 and NAb binding and classify the direction and magnitude of epistatic interactions between variant mutations and the broader mutational repertoire. The results reveal only weak epistasis which, although more pronounced for the Gamma and Omicron variants than for the Delta variant, suggests limited potential for continued evolution towards more complete vaccine escape.

## Results

### Rationale

Epistatic interactions among mutations in the RBD of SARS-CoV-2 S protein are of interest and concern because they might substantially increase the risk of vaccine escape. Mutations in the RBD subtly change the shapes of the interfaces between RBD and ACE2, and between RBD and NAb (Figure 1A). While the structure of the WT RBD-ACE2 interface is highly similar to that of the RBD-NAb interface (see below), a single mutation in the RBD can result in distinct shape changes in both interfaces. These changes can be depicted by the position of each mutant on the plane bounded by the receptor binding cost and the antibody cost (Figure 1B). The cost is the increase (positive cost) or decrease (negative cost) in the ACE2 or NAb binding affinity relative to the WT. The four quadrants of this plane (Figure 1B) represent four broad categories of mutations. Mutations in the top, right quadrant are strongly destabilizing relative to both ACE2 and the NAb. The bottom, right quadrant contains mutants that strongly destabilize the interaction with ACE2 but not with the NAb. Most mutants in these quadrants are not evolutionarily viable. Mutants in the bottom, left quadrant stabilize or only weakly destabilize both interfaces. These mutations may or may not provide a selective advantage to the virus depending on the fraction of the host population that has been vaccinated or has recovered from prior infection. The top, left quadrant contains mutations that strongly destabilize the interaction with NAb but not with ACE2 and therefore are most likely to admit vaccine escape.

**Figure 1.**
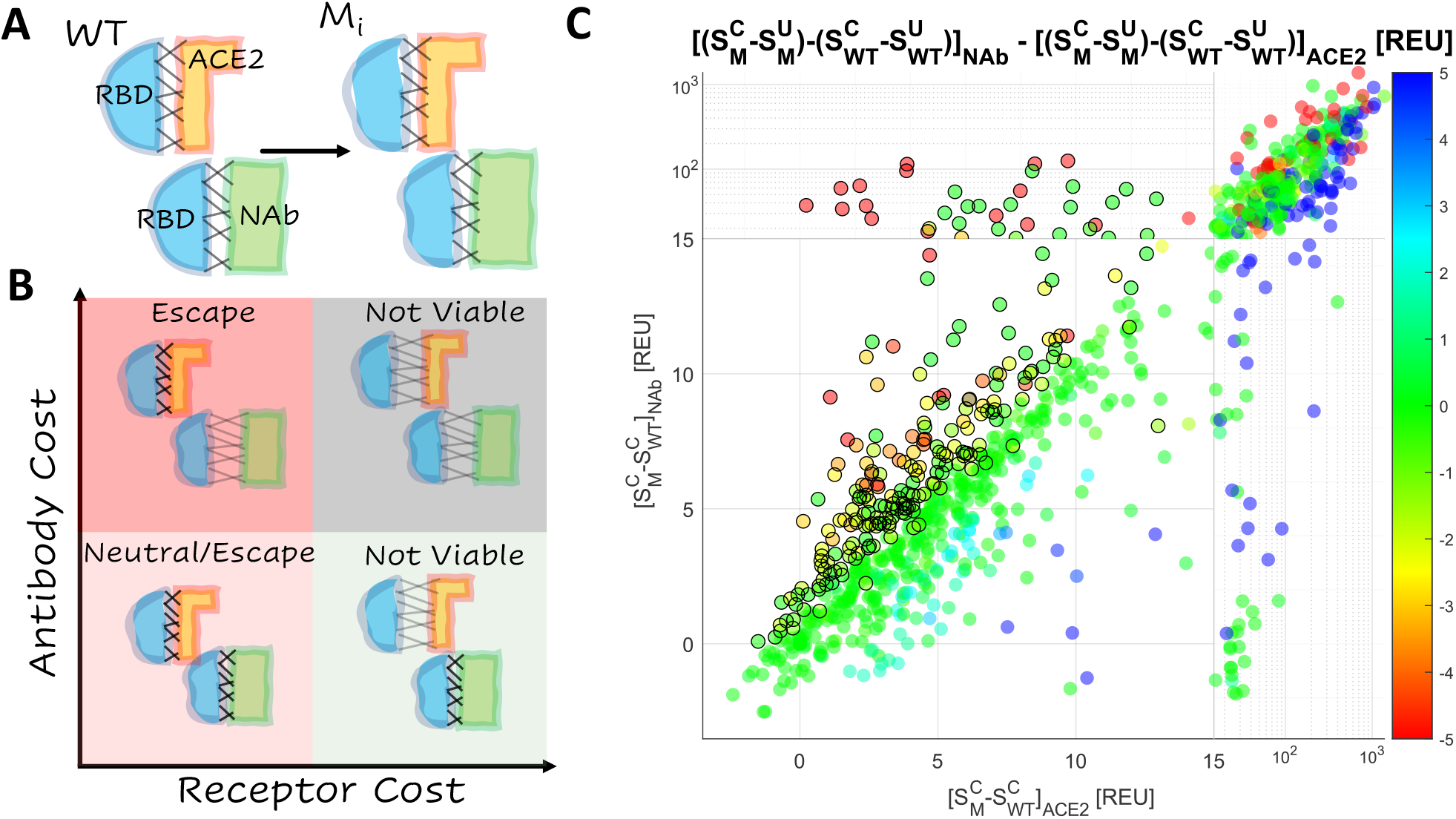
Landscape of Vaccine Escape Mutants for the WT RBD. **A**. Cartoon depicting unique conformational changes to the RBD (blue) in complex with ACE2 (orange) and the NAb (green) associated with the same mutation. **B**. Cartoon depicting the landscape of vaccine escape mutations (the plane of receptor cost vs antibody cost). **C**. Landscape of vaccine escape mutations for the WT RBD. Circles with a black outline are NAb escape candidates. Color indicates propensity for escape as measured by *ΔΔG*.

### Single-mutant vaccine escape candidates for the wild type RBD

Starting with the two crystal structures of interest, the RBD in complex with ACE2 https://www.rcsb.org/structure/6M0J(16) and the RBD in complex with the NAb CV30, https://www.rcsb.org/structure/6XE1, we generated a representative ensemble of 50 native conformations per complex following standard Rosetta protocols (see Methods and Discussion for details). Although NAb that bind epitopes, which do not overlap with the RBD have been identified(8), at the time of writing, the antibodies most critical for assessing the risk of vaccine escape appear to overlap with the RBD(9) and are well represented by CV30. Regions important for antibody binding are known to overlap broadly among human coronaviruses(17). We then identified the RBD residues at the interface for each conformation and, in all conformations, introduced all single amino acid substitutions at these sites. For the WT and Delta variant, 52 residues (Table 1) were identified at the interface of at least one conformation for either complex. Four additional residues were identified for the Gamma and/or Omicron variants. Sites 480 and 488 are connected by a disulfide bond and were found to be unsuitable for substitution.

**Table 1:** Receptor-binding and antibody-binding interface footprints in the RBD.

|  |  |
| --- | --- |
| WT RBD-ACE2 Footprint | 403, 405, 417, 445-447, 449, 453, 455-456, 473-478, 484-491, 493-498, 500-506 |
| Delta RBD-ACE2 Footprint | Same as WT |
| Gamma RBD-ACE2 Footprint | WT + 404, 406, 408, 439, 499 - 484 |
| Omicron RBD-ACE2 Footprint | WT + 404, 439, 499 - 445, 484, 491 |
| WT RBD-NAb Footprint | 403-406, 409, 414-417, 419-421, 446-447, 449, 453, 455-461, 473-478, 484-498, 500-506 |
| Delta RBD-NAb Footprint | Same as WT |
| Gamma RBD-NAb Footprint | WT + 408, 480 |
| Omicron RBD-NAb Footprint | WT - 446 |

All 19 substitutions in each of the remaining 54 sites were investigated, with the exception of WT reversion for the variants. Note that substitutions made in the WT are all single mutants; those in the Delta variant are triple mutants (given that this variant contains two mutations in the RBD); those in the Gamma variant are quadruple mutants; and those in the Omicron variant each encompass 16 mutations in total. We chose to examine all variants containing Delta/Gamma/Omicron substitutions and an additional substitution at the interface rather than probing a randomly selected, representative ensemble of multiple mutants relative to the WT because the mutations present in the Delta/Gamma/Omicron variants are known to be of biological and epidemiological relevance and the space of all such multiple mutants is prohibitively large. Altogether, this analysis produced more than 400k structures which necessitated the development of a computationally efficient, and therefore simplified, protocol. To address this need, mutants were introduced into each conformation without repacking of adjacent sidechains or backbone minimization. This minimalist approach yielded favorable comparisons to the available experimental data (see below). However, generally, substitutions might introduce steric or charge clashes within the conformations in which the mutations were introduced (without repacking and minimization). Inference of the relative change in binding affinity for the ACE2 and NAb complexes is limited for such mutations. However, we observed a favorable comparison to experimental data in this respect as well, whereby few experimentally predicted escape mutations (relative to the WT) fall into this, inference-limited, category (see below).

Structure stability was estimated by the total score, *S*, in arbitrary units produced by the empirically-driven Rosetta Energy Function 2015(18) (labelled REU for “Rosetta Energy Units”, https://new.rosettacommons.org/docs/latest/rosetta_basics/Units-in-Rosetta). The total score was calculated for each of the 50 conformations of the NAb and ACE2 complexes generated, and the mean value was assessed with, 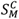, and without, 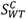, the mutation. The receptor cost and antibody cost were estimated as 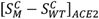 and 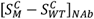, respectively. The interface free energy (*ΔG*) was also more directly approximated by the difference between the total score of the unbound state and the complex, *S*^*C*^ − *S*^*U*^. The effect of the mutation on this value (*ΔΔG*) was reported for both complexes,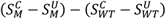. Figure 1C shows the distribution of RBD mutations on the plane bounded by the ACE2 and NAb binding costs and putative Nab escape candidates, for which 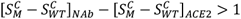 or *ΔΔG*_*NAb*_*-ΔΔG*_*ACE2*_*>1* and 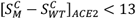. The threshold value of 13 was selected to remove from consideration mutations that likely produce steric or charge clashes in the structure; few experimentally validated escape candidates were observed above this value (see below).

Mutants showed strong clustering along the diagonal (identity line: receptor cost is equal to antibody cost), indicating that most mutations similarly affected the WT RBD-ACE2 and RBD-NAb complexes. Mutations in the top, left quadrant of the plane, which corresponds to strong destabilization of the interaction with NAb but not with ACE2, are the strongest candidates for vaccine escape, followed by those in the bottom, left quadrant, which includes weakly destabilizing mutations. The selective advantage (or lack thereof) of mutations in this quadrant depends on the fraction of the host population that has been vaccinated or has recovered from prior infection (see Discussion). In a fully vaccinated population, mutations that substantially reduce infectivity through the destabilization of receptor binding, while also destabilizing the interaction with NAb, could still provide a selective advantage. In particular, multiple mutations in 6 sites (417, 477, 484, 491, 493, 499) were found to substantially destabilize the RBD-NAb complex relative to the RBD-ACE2 complex (Figure S1). Additionally, we identified site 453 to harbor mutations that simultaneously stabilize the RBD-ACE2 complex and destabilize the RBD-NAb complex. These observations are broadly consistent with the results of deep mutational scanning(8, 9, 19-22). Therefore, we conservatively considered all mutations, for which the antibody cost exceeded the receptor cost, to be viable escape candidates.

### Single-mutant vaccine escape candidates for the Gamma, Delta, and Omicron variant RBDs and predicted epistatic interactions

Having charted the ACE2-binding and NAb-binding landscapes for the WT RBD, we sought to identify the most prominent combinations of RBD mutations circulating over the course of the pandemic. As of June, 16^th^, 2021, there were 53 countries, from which more than 1,000 SARS-CoV-2 isolates were contributed to the GISAID(23) database (see Data Availability). For each of these locations, we randomly selected 1,000 isolates and reported the frequency of each combination of RBD mutations among the 53,000 selected isolates over time. Figure 2A displays this region-normalized global prevalence of the 10 most common combinations of RBD mutations. The 6 RBD single-mutants (501Y/Alpha Variant, 477N, 439K, 484K, 478K, and 459F) began emerging between July and November 2020. All these single mutants were eventually displaced by 4 RBD multiple mutants (452R|478K/Delta Variant, 417T|484K|501Y/Gamma Variant, 417N|484K|501Y/Beta Variant, and 346K|484K|501Y), which began emerging in November, 2020, with the exception of Beta, which according to our analysis, first appeared in July. By March, 2021, the WT had become less prevalent than the Alpha, Gamma, and Delta variants. We pursued further analysis for the Delta, Gamma, and the recently identified Omicron variant RBDs given their high and rising global prevalence. The complexes were prepared starting from the WT crystal structures and treated identically to the WT. We also discuss how similar results may be obtained starting directly with the native conformations approximated for the WT, substantially reducing the computational burden (see Methods).

**Figure 2.**
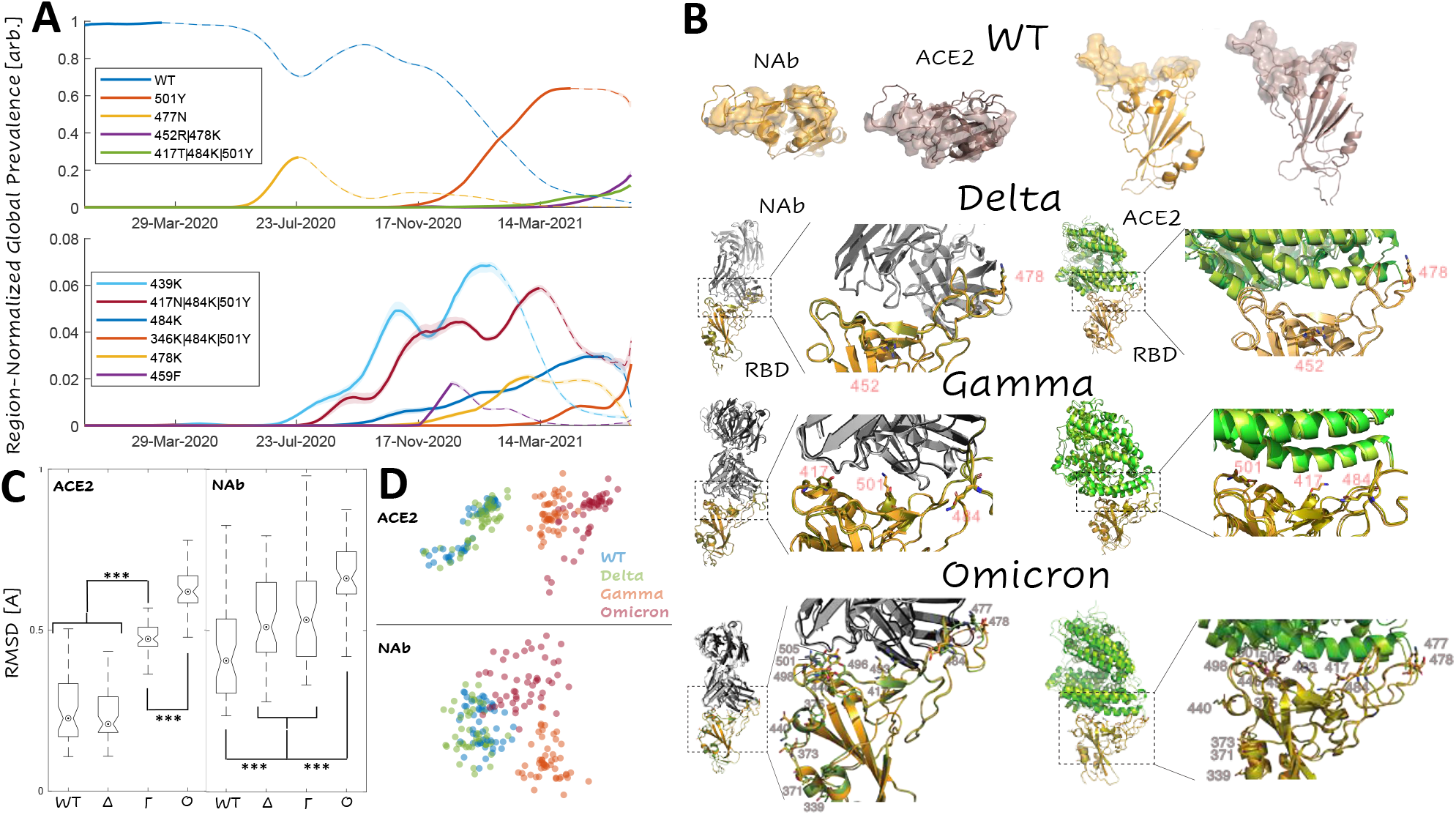
Dominant Trends in Circulating RBD Mutations. **A**. Region-normalized global prevalence of the top 10 most common combinations of RBD mutations over time. Lines are solid up to peak prevalence and dashed afterwards. Shading indicates confidence intervals. **B**. Structural comparison of the complexes of ACE2 and NAb with RBD for WT, Delta, Gamma, and Omicron variants. *Top*: WT footprints including the residues with an interaction within 4A of the partner. *Bottom:* Visualization of the Delta, Gamma, and Omicron variant interfaces. Mutations are labeled and represented as sticks. WT structures are superimposed for the RBD of each variant: WT, orange; variant, olive **C**. Interface RMSD for ACE2 and NAb complexes relative to an arbitrary WT conformation (over spike protein sites 403-406,408-409,414-417,419-421,439,445-447,449,453,455-461,473-478,480,484-506). Asterisks denote p-values less than 0.02 for a Wilcoxon Rank Sum Test. **D**. CMDS applied to the pairwise RMSDs among all RBD-ACE2 and RBD-NAb complexes. 2D visualizations of 3D projections are displayed (see Figure S2 for 3D).

The rapid emergence and subsequent displacement of RBD single mutants might in part result from epistasis among the RBD variant residues or could be due to purely additive interactions. The most prominent trend was the displacement of the single mutant, 501Y (Alpha variant) by the Beta and Gamma variants (both also containing 501Y). Residue 501Y has been shown to substantially increase the binding affinity with ACE2 which, however, is reduced with the addition of mutation 417N in the Beta variant(11). In contrast, 417N severely reduces the neutralizing activity of a variety of NAb(24). These observations imply that mutations in site 417 provide a selective advantage through destabilization of the NAb complex, but given the large overlap between the RBD-NAb and RBD-ACE2 interfaces, maintenance of sufficient infectivity requires a compensatory mutation, such as 501Y, that stabilizes the RBD-ACE2 complex. Note that the emergence of these variants preceded widespread vaccination and that, although the competition between antibody and receptor binding is present even during an infection of a naïve host, evolutionary pressures are likely to shift with increasing rates of vaccination and prior infection (see below). At the time of writing, the origin of the Omicron variant, and thus the evolutionary pressures that led to its emergence remain unknown (see discussion); however, despite bearing many more RBD mutations, the epistatic landscape at the interface is highly similar between the Gamma and Omicron variants (see below).

Examination of the interface footprints, defined as the ensemble of sites predicted to lie at the interface of at least one of the 50 conformations for each complex, for the 8 complexes of interest, demonstrates that the RBD makes a greater number of contacts with NAb than with ACE2 within the same range of sites, 403-506 (Figure 2B). The footprints of the WT and Delta variant interfaces in both the RBD-ACE2 and the RBD-NAb complexes are identical. The WT/Delta RBD-ACE2 footprint consists of 37 sites whereas the WT/Delta RBD-NAb footprint includes 51 sites (Table 1). The Gamma RBD-ACE2 footprint consists of 41 sites including all those in the WT/Delta footprint, with the single exception of site 484, and five additional sites. The Gamma RBD-NAb footprint consists of 53 sites including all those in the WT/Delta interface and, in addition, sites 408 and 480. The Omicron RBD-ACE2 footprint consists of 37 sites with three WT sites missing (including 484) and three additions. The Omicron RBD-NAb footprint includes 50 sites with one WT site missing.

Sites in the RBD-NAb footprint that are not shared by the RBD-ACE2 footprint might provide routes for the emergence of vaccine escape variants. However, because the RBD-ACE2 footprint is smaller than the RBD-NAb interface, the former is more sensitive to perturbation than the latter, for example, from mutations in site 417, which is part of the footprint of all 8 complexes. Notably, site 484 is absent from the Gamma and Omicron RBD-ACE2 footprints, but remains in the respective RBD-NAb footprints. Also of note, site 446, which is mutated in Omicron, is present in the RBD-NAb footprint but not the RBD-ACE2 footprint for this variant. Consistent with the differences in the footprints, we found the Omicron and, to a lesser extent, Gamma variant RBD conformations in complex with ACE2 to be significantly different from that of the WT and Delta variants, which could not be differentiated from one another. All variant RBD conformations in complex with the NAb were found to be significantly different from that of the WT. For Delta and Gamma, this difference was modest and smaller in magnitude than the variability among the RBD-NAb conformations, whereas Omicron showed a more pronounced difference from the WT (Figure 2C).

Classical multidimensional scaling (CMDS) applied to the pairwise interface RMSDs among all RBD-ACE2 and RBD-NAb complexes showed that the Gamma and Omicron RBD-ACE2 structures lie on a shared continuum of conformational change relative to the WT, with the Omicron conformations being closer to Gamma than to the WT. In contrast, Omicron, Gamma, and to a lesser extent, Delta RBD-NAb structures all represent distinct conformational changes relative to the WT (Figures 2D and S2). Despite these differences, the effects of mutations in the RBD were found to be principally additive in all variants, that is, there seems to be little epistasis.

When a second mutation, *M*_*j*_, is introduced in addition to a prior mutation, *M*_*i*_ (Figure 3A), the resulting conformational change can be additive so that the effect of the two mutations is the sum of the effects of the two individual mutations. In this case, the position of the double-mutant *M*_*i,j*_ on the plane defined by the receptor cost and antibody cost relative to the single mutant, *M*_*i*_, will be the same as that of the single-mutant, *M*_*j*_, relative to the WT. If the conformational change is non-additive, representing an epistatic relationship, the resulting trends can be classified by their impact on potential vaccine escape. Such trends could be escape-neutral when the ensemble of candidate vaccine escape mutations differs from that for the WT, but the number of such candidates is the same; escape-minimizing when the antibody cost is on average reduced relative to the receptor cost across all mutations for the mutant vs the WT; or escape-exacerbating where the antibody cost is on average increased.

**Figure 3.**
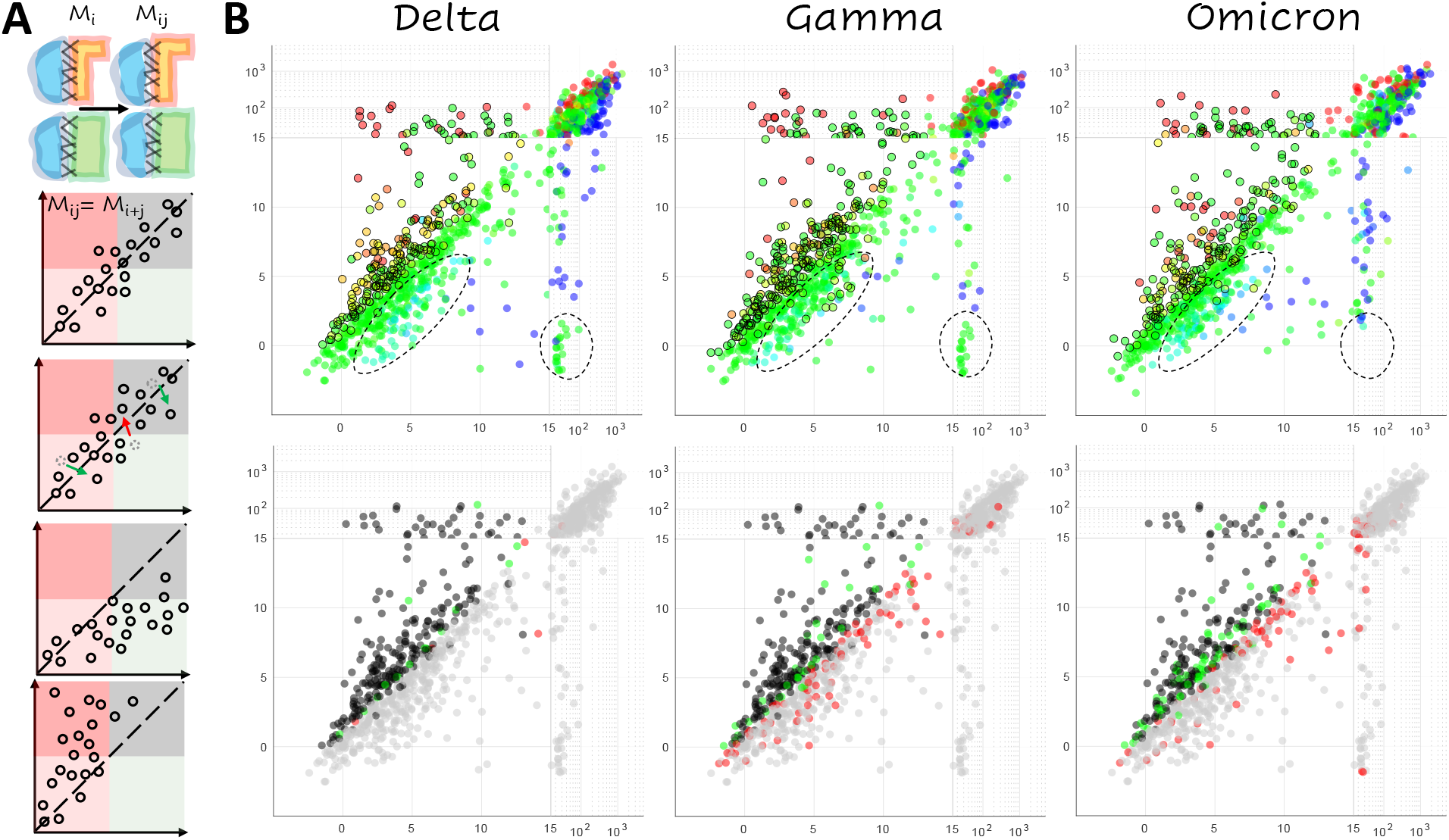
Epistasis within the RBD. **A**. Cartoon illustrating additive and non-additive (epistatic) interactions between mutations. From top to bottom: additive, escape-neutral, escape-minimizing, and escape-exacerbating. *Mi, Mj, Mij* denote the effects of the single and double mutants. **B**. *Top:* Landscape of vaccine escape mutations for the variant RBDs. Coloring, as in Figure 1C, indicates propensity for escape as measured by *ΔΔG*. Circles with a black outline denote NAb escape candidates. Dashed lines highlight differences among variants. *Bottom:* Landscape of vaccine escape mutations for the WT RBD. Black points are candidates for both WT and variant; gray points are not candidates for either WT or variant; green points are only candidates for WT; red points are only candidates for the variant.

The landscape of mutants predicted to enhance vaccine escape for the Delta variant was almost identical to that of the WT but differed significantly from the Gamma and Omicron landscapes (Figure 3B). These trends are summarized in Table 2, which tabulates all non-shared candidates. There are 15(13) escape candidates in the WT that were not predicted to enhance escape for Delta and 6(2) candidates in Delta but not WT (values in parentheses are mutations with 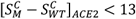, regardless of whether or not the mutation is a candidate, a threshold chosen to mitigate potential artifacts caused by steric or charge clashes). In contrast, in the case of Gamma, there were 32(28) candidates identified in WT but not Gamma, and 86(66) candidates identified in Gamma but not the WT. Omicron demonstrated intermediate behavior with 67(59) candidates identified in WT but not Omicron, and 75(48) candidates identified in Omicron but not the WT. Thus, we identified 9(11) fewer escape candidates for Delta compared to the WT, but 54(38) additional candidates for Gamma, and either 8 additional candidates or 11 fewer candidates (ignoring potential steric/charge clashes in the WT) for Omicron.

**Table 2:** Numbers of antibody-escape candidates.

|  | Total |  | WT not Variant | Variant not WT | Difference |
| --- | --- | --- | --- | --- | --- |
| WT | 241 | $\Delta$ | 15(13) | 6(2) | -9(-11) |
| Delta | 232 | $\Gamma$ | 32(28) | 86(66) | 54(38) |
| Gamma | 295 | O | 67(59) | 75(48) | 8(-11) |
| Omicron | 249 |  |  |  |  |

Values in parentheses are tabulated mutations with $[S_M^C - S_{WT}^C]_{ACE2} < 13$ , for both the WT and variant.

Consistent with the more dramatic conformational change observed in the RBD-ACE2 complex relative to the RBD-NAb complex, the non-additive effects observed in the Gamma variant appear to predominantly result from the decreased sensitivity of the RBD-ACE2 interface to mutation. This conclusion is compatible with the available experimental results. Figure S3 shows the distribution of the receptor cost, 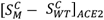, for two categories of mutations and for each of the three receptor complexes: those at the interface that have been experimentally demonstrated to reduce neutralizing activity of antibodies COV2-2050 and COV2-2479 in the WT(8), and all others (included in the same experimental study) at the interface. As discussed above, the upper bound for the receptor cost, 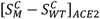, is lower for mutations predicted to reduce NAb activity than for other mutations. However, the Gamma candidate ensemble exhibits a reduced median receptor cost. In other words, in the Gamma variant, mutations that are predicted to reduce NAb activity are also less likely than other mutations to reduce the receptor binding affinity relative to the WT and Delta variant. The three residues within the RBD-ACE2 footprint of the Omicron variant not present in the WT overlap with those of the Gamma variant (404, 439, and 499) and, although not statistically significant, the decreased cost of receptor binding was observed for Omicron as well. The Omicron variant additionally displayed a modest reduction in median receptor cost for the broader category of experimentally studied mutations, suggesting greater flexibility at the receptor interface (Figure S3).

Figure 4 summarizes the magnitude of the increased risk of vaccine-escape for each mutation at the RBD interface (given 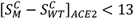) for each variant relative to the WT. Epistasis increasing the risk of vaccine-escape is primarily apparent in three regions of the RBD interface: site 417, site 477, and site 494 together with the surrounding neighborhood. The trend in site 417 was observed only in the Gamma and Omicron variants, which already contain mutations at that site, showing that further changes to this site could result in enhanced vaccine escape. However, the epidemiological implications of this finding are limited considering that mutations in site 417 are likely to pose a risk of vaccine escape in most variants. The enhanced escape associated with mutations in site 477 for all variants relative to the WT, together with the early spread of 477N and the presence of 477N in the Omicron variant, suggest that this site could play an important role in host adaptation. Most prominently, mutations in site 494 and the surrounding neighborhood are likely to enhance vaccine escape in all variants. Indeed, 494P has both been found in circulation and experimentally demonstrated to reduce antibody neutralization capacity of convalescent sera(25).

**Figure 4.**
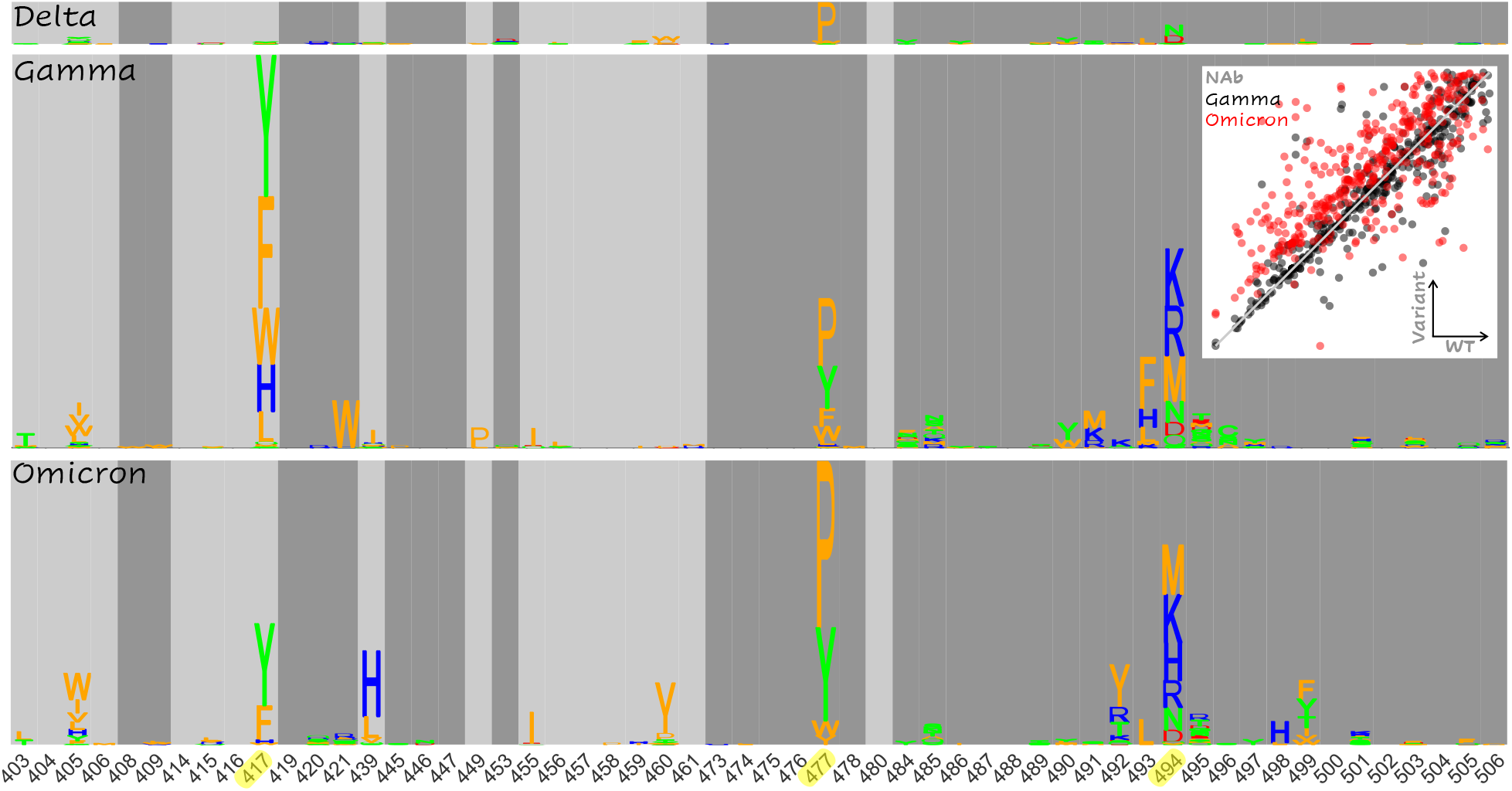
Landscape of Epistatic Effects Supporting Enhanced Vaccine Escape. Non-additive escape-exacerbating motifs in Delta (top), Gamma (middle), and Omicron (bottom) variants. The size of each letter corresponds to the increased likelihood of vaccine escape for the substitution in the variant relative to the WT. *Inset*: Total score change (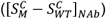) induced by mutation in the Gamma (black) and Omicron variants (red) vs the WT (identity line overlaid). The minimum value (less 1) is subtracted from each distribution for normalization (see Figure S5 for more).

In addition to these apparent differences among the ensembles of candidate vaccine-escape mutations, we observed sites that harbored no candidates, but nevertheless displayed signatures of increased risk of vaccine escape for Gamma and/or Omicron. The two most notable trends were observed in sites 408 and 504 (Figure S4). All but one substitution in site 408 enhance vaccine-escape for Gamma and Omicron, but surprisingly, all have the opposite effect in Delta. Similarly, all substitutions at site 504 substantially enhance vaccine-escape in Gamma and Omicron but exert a modest opposite effect in Delta. However, these mutations are not considered candidates in our analysis because, even for Gamma and Omicron, they destabilize the ACE2 interaction to a greater extent than the interaction with NAb. Additionally, 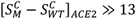 for substitutions at site 504, which limits confidence in the assessment of trends at this site.

Importantly, few differences were observed between the epistatic landscapes of the Gamma and Omicron variants, mainly in sites 417, 439, 499 (see Figure 4), and 446 (see Figure S4). Sites 417 (reduced escape-related epistasis) and 446 (increased escape-related epistasis) are already mutated in Omicron and the signatures in sites 439 and 499 are modest relative to the consistent trend observed in site 494. Nonetheless, it is important to acknowledge a small, systematic bias towards NAb destabilization within the Omicron variant that is not observed within Gamma or Delta (Figure 4, inset and Figure S5). As demonstrated above, unlike the RBD-ACE2 complex, which displays a shared continuum of conformational change relative to the WT, RBD-NAb structures all represent distinct conformational changes (Figures 2D and S2) with the greatest change observed for Omicron. Most mutations, which modestly destabilize the WT RBD-NAb complex (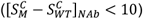), are slightly more destabilizing in the Omicron RBD-NAb complex compared to the other variants. As discussed above, only a subset of these mutations are considered candidates for antibody escape, which depends on the properties of the mutation within the RBD-ACE2 complex as well. The principal escape signatures are conserved between the Gamma and Omicron variants, with fewer escape candidates identified overall for the Omicron variant. However, this systematic destabilization of the Omicron RBD-NAb complex suggests, in principle, additional avenues towards vaccine escape.

## Discussion

Here we report the results of a computational study predicting the effects of all single mutants at the RBD-NAb and RBD-ACE2 interfaces for the WT as well as the Delta, Gamma, and Omicron variants of SARS-CoV-2 on receptor and antibody binding. For the WT, we found multiple mutations in 6 sites (417, 477, 484, 491, 493, 499) that are predicted to significantly destabilize the RBD-NAb complex relative to the RBD-ACE2 complex and appear to pose a risk of vaccine escape, which is broadly consistent with the results of deep mutational scanning(8, 9, 19-22). Overall, most mutations at the interface were found to similarly effect the WT and all variants, indicating limited epistasis at the interface. Non-additive, epistatic interactions predicted to increase the risk of vaccine-escape were apparent, however, at sites 477 and 494 as well as in the surrounding neighborhood. This trend is particularly prominent in the Gamma variant so that, across all sites at the interface, we predicted 22% more escape candidate mutations for the Gamma variant than for the WT. In contrast, there is little apparent epistasis in the Delta variant, and across all sites at the interface, we predicted 4% fewer candidate mutations compared to WT. Despite harboring many more RBD mutations and displaying a small, systematic trend towards NAb complex destabilization, the Omicron variant demonstrated intermediate behavior with only 3% more candidate escape mutations compared to the WT.

Epistasis is a major if not the principal driver of protein evolution(26). Compensatory mutations are particularly strong epistatic interactions that can result in chemotherapeutic(27) or antimicrobial(28) drug resistance, and are commonly observed throughout species evolution(29). In a completely susceptible population, mutation 501Y, which appears to substantially increase infectivity(11), is expected to evolve under positive selection. As a population gains immunity, through prior exposure and/or vaccination, selective pressures rapidly change to promote the emergence of resistant variants(7). Under these conditions, 501Y and other mutations, which increase infectivity might primarily play the role of compensators for mutations destabilizing NAb interactions, such as 417N or 417T. As global vaccinations rise, due to changing evolutionary pressures, it can be expected that more mutations emerge that destabilize the interactions of the RBD with both NAb and ACE2, thus resulting in (partial) escape variants that, however, also have reduced infectivity. However, variants such as Gamma that carry both an antibody destabilizing mutation and a compensatory mutation have the potential to undercut this trend.

At the time of writing, the origins of the Omicron variant are unclear and is suspected to have evolved over an extended period of time within an immunocompromised individual(s) or an animal reservoir(30). Such environments could present selective pressures distinct from the broader human host population, but the changing competition between receptor and antibody binding described above is likely to be conserved. Despite the more pronounced conformational change, the average behavior of the conformational ensemble selected to represent the Omicron variant was found to be less destabilizing for the receptor compared to Gamma but highly destabilizing for the antibody, indicating that our model for Omicron is likely to be at least as accurate if not more accurate than those for the Gamma and Delta variants (Figures S12-13).

Above all else, we find it important to highlight three points. 1) The epistatic landscapes of the Gamma and Omicron variants are highly similar (Figure 4). 2) We find fewer, not more, escape-exacerbating mutations in Omicron compared to Gamma (Table 2). 3) These features persist in spite of the fact that RBD-ACE2 structures represent a shared continuum of conformational change relative to the WT, with the Omicron conformations being closer to Gamma than the WT (Figures 2D and S2), indicating that the magnitude of conformational change does not trivially correlate with the propensity for exacerbated antibody escape. The neutralizing activity of existing vaccine elicited NAb against the Omicron variant is likely to be substantially reduced(31), but multi-dose vaccination is expected to recover efficacy(32). Our results emphasize that, although the adaptive repertoire of SARS-CoV-2 may be robust, structural constraints on the RBD make continued evolution towards more complete vaccine escape unlikely, suggesting continued efficacy of the existing vaccines.

The work presented here is strictly computational, and although we demonstrate agreement with experimental results where possible, many features not captured by the models presented (involving protein expression, docking, and other factors) could modulate antigen-receptor and/or antigen-antibody binding. Furthermore, although we explore many conformations for both the RBD-ACE2 and RBD-NAb interfaces, we start from a single crystal structure for each. We believe the conformational ensembles selected to represent each complex are diverse enough to accurately reflect the relative destabilization of the NAb and ACE2 complexes across the spectrum of RBD interface mutations, which is the primary concern. However, this conformational diversity makes it difficult to demonstrate stabilizing interactions, which are typically much weaker than destabilizing ones(12). Although multiple low-energy conformations were resolved for all variants and the WT, the average behavior of the conformational ensemble selected to represent the Delta variant relative to the WT was found to be weakly destabilizing for NAb and neutral for ACE2, whereas in the case of Gamma, it was weakly destabilizing for both complexes. This is unlikely to accurately reflect the relative binding affinities between these variants and the WT given the enhanced infectivity of both variants, particularly Delta(14). However, it should be recognized that the relationship between the measures of interface stability we report and viral life history traits (infectivity, immune activity, etc.) is complex.

Although we believe we proposed sensible thresholds for determining which structures can be analyzed with high confidence and the biological implications of the relative destabilization of the NAb vs ACE2 are overt, the effects studied in this work do not represent the entire diversity of possible host adaptation. It is incompletely understood at the time of writing why the Delta variant appears to replicate faster than the WT(14) and substitutions outside the Spike protein could play key roles in immune modulation(33, 34). Even less is known about the Omicron variant. A targeted exploration of the lowest-energy conformations achievable for each variant might yield better agreement with the known properties of these variants and, in particular, reveal stabilizing RBD-ACE2 interactions. However, this would likely come at the cost of generalizability and decrease the power of our approach to predict relative destabilization of interface mutations between NAb and ACE2 complexes.

We also emphasize that we limit our study to the RBD of the S protein. In principle, epitopes located outside the RBD and far from the interface could play an important role in the emergence of vaccine escape variants by decreasing the similarity between the interaction with the antigen and receptor, and that between the antigen and an antibody(35). However, if such effects were dominant, the satisfactory agreement between our predictions, deep mutational scanning, and the observed frequency of mutations within the RBD among the circulating variants, presumably, would not be recovered. Nevertheless, many routes of adaptive evolution are potentially available to this virus so that agreement with prior results is not a guarantee of predictive success. Although we believe our results strongly suggest a limited repertoire of escape-mediating mutations within the RBD, the possibility should be considered that mutations outside the RBD have the potential to increase this repertoire.

Finally, we note that the space available for neutral evolution, even within the RBD alone, is large. In principle, this makes possible the acquisition of many RBD mutations, some combinations of which might exhibit substantially greater escape-exacerbating epistatic effects than the variant substitutions explored in this work. However, this appears unlikely considering the large number of mutations observed in the Omicron variant, 15. We believe this variant is an excellent test case to probe the limits of epistatic potential at the RBD interface.

## Conclusions

We employed a computational approach to study the effects of all single mutations at the RBD-NAb and RBD-ACE2 interfaces for the WT as well as the Delta, Gamma, and Omicron variants of SARS-CoV-2. Overall, little epistasis at the RBD interface was detected, with additive effects on the binding affinities observed for most pairs of mutations. In the Delta variant, the observed non-additive trends weakly stabilize the interaction of the RBD with the NAb relative to the interaction with ACE2, whereas in the Gamma variant, epistasis is predicted to more substantially destabilize interaction with the NAb relative to ACE2. The epistatic landscape of the Omicron variant closely resembles that of Gamma, with an additional small systematic bias towards NAb destabilization, but with fewer predicted escape candidates overall. These results suggest that, although the Omicron variant poses new risks not observed for Delta, including the evolution towards greater NAb destabilization, structural constraints on the RBD make the continued evolution towards more complete antibody escape unlikely.

The modest ensemble of mutations relative to the WT that are currently known to reduce vaccine efficacy is likely to comprise the majority of all possible escape mutations for future variants, predicting continued efficacy of the existing vaccines.

## Methods

### Selection of Crystal Structures

In this work, we considered a single crystal structure of the SARS-CoV-2 spike protein Receptor Binding Domain (RBD) in complex with the human receptor, Angiotensin Converting Enzyme 2 (ACE2), PDB:6M0J(16). While there are likely multiple mutations outside of the RBD which significantly affect binding characteristics of the spike protein, as has been demonstrated for site 614(36), the structure of the RBD itself is unlikely to be substantially modified by such mutations. Thus, while only able to reveal a subset of mutations of interest, the focused study of RBD complexes presented here remains biologically realistic.

Similarly, we consider a single crystal structure of the RBD in complex with a neutralizing antibody (NAb),CV30 PDB:6XE1(37). CV30 was recognized early on as the most potent NAb observed within the sera of a SARS-CoV-2 positive donor while the majority of antibodies were found to target non-neutralizing epitopes outside of the RBD(38). Subsequently, it became clear that antibodies targeting epitopes outside the RBD, particularly those targeting the N-terminal domain (NTD), likely play a protective role and mutations reducing the affinity of these antibodies may be epidemiologically significant(35, 39). As acknowledged above regarding the selection of the spike crystal structure, while our focus on CV30 is only able to reveal a subset of mutations of interest, the mutations discussed in this work predicted to affect CV30 binding affinity are likely highly epidemiologically relevant.

### Construction of Representative Ensembles of Interface Conformations

Protein crystal structures may differ substantially from the native conformations(40). Throughout this work, we utilize the Rosetta(13) software suite to approximate both wild type (WT) and mutant conformations of the receptor and NAb complexes. All protocols used throughout were implemented using the RosettaScripts(41) package and the XML files used along with the associated executed command lines are made available in Table S1. Approximation of the native conformational ensemble may be separated into two steps, identifying the optimal side chain conformation (repacking) and moving the protein backbone (minimization), to minimize the energy function applied. This may be accomplished using the Rosetta Relax application which iteratively applies each of these two steps.

Beginning with the crystal structure, we iteratively applied the FastRelaxMover using default parameters (with the exception of disabling design) for up 12 iterations(15 for Omicron) and up to 1000 repeats. We found the total score to be insensitive to additional applications of FastRelax after 5 iterations on average. Each resulting structure was scored using the InterfaceAnalyzerMover, repacking the unbound state but not the bound state (as the input complex has already been optimized).

This protocol returns the total score, *S*, in arbitrary units produced by the empirically-driven Rosetta Energy Function 2015(18) (labelled REU for “Rosetta Energy Units”, https://new.rosettacommons.org/docs/latest/rosetta_basics/Units-in-Rosetta), as well as dG separated which is the difference in the total score between the bound (complex) and unbound state, *S*^*C*^*-S*^*U*^, derived from separating the binding partners. This protocol may also be used to identify the residues within the complex which constitute the interface.

Such an ensemble of structures often forms an “energy funnel”(42) where the root mean square distance (RMSD) between superimposed backbone carbon atoms of each structure and the structure with the lowest total score or the lowest dG separated is positively correlated with the total score of that structure. In order to evaluate whether such a funnel exists for these ensembles, we identified the structure with the lowest dG separated and 90 residues predicted to be interface residues within the RBD in at least one conformation in addition to the (+/-) 3 adjacent amino acid neighborhood of each such residue (sites 400-424, 440-464, and 470-509 in the spike protein for WT; additional sites 436-439 for Delta; additional sites 434-439 for Gamma; additional sites 434-439 and 510 for Omicron).

Figure S6 displays dG separated vs the interface RMSD for both complexes for the WT. Few structures appear in the lower left corner of each plot from only one or two (NAb or ACE2 respectively) independent “trajectories” of iterative FastRelax application beginning with the crystal structure. Furthermore, while the lowest dG separated is more than 30% greater in magnitude than the highest for both complexes, the interface RMSD between any complex and the minimum dG separated complex is less than an angstrom. These findings suggest selecting the single minimum dG separated conformation for either complex is unlikely to constitute a realistic model of the native interface and may in fact represent an unrealistic, entropically disfavored state(43).

Figures S7-9 display dG separated vs the interface RMSD for both complexes for the Delta, Gamma, and Omicron variants. The distribution of conformations for the NAb is similar to the WT for both Delta and Gamma; however, the minimum dG separated obtained for Gamma is higher than that for the WT suggestive of antibody destabilization for this variant. dG separated for Omicron is substantially higher indicating significant destabilization. For Delta and Omicron, the distribution of conformations for ACE2 more resembles a funnel suggestive of receptor stabilization for these variants. For Gamma, there appear to be 2-3 distinct, equally low energy conformations for ACE2 which makes the interpretation of the energy landscape more challenging. For consistency, representative conformations were selected according to the same protocol for all variants.

The construction of an ensemble of representative conformations is desired so that the average behavior of such an ensemble is likely to reflect that of the native complex. Ideally all available structures would be statistically weighted and included in this ensemble; however, this is computationally intractable. Instead we selected 50 conformations for each complex as follows (see Figures S10-13): 1) The conformation corresponding to the minimum total score after any iteration of FastRelax from each independent “trajectory” beginning with the crystal structure was selected. 2) The 10 structures with the lowest total score were removed and the 10 structures with the lowest dG separated were removed (these are not identical structures and as discussed above, may represent unrealistic, entropically disfavored conformations). 3) The remaining structures were ranked by total score, *r*_*S*_, and dG separated, *r*_*G*_, and the 50 structures with the lowest composite rank, *r*_*S*_*+r*_*G*_, were selected. The PDB files for these structures are made available (see Data Availability). We believe the equally-weighted average behavior of these conformations, which are stable with well-resolved interfaces, constitute a reasonable model of the native complexes.

### Analysis of Single Mutants Relative to WT, Delta, and Gamma

Given the 100 structures selected as described above (50 for each complex), a more restrictive list of 52 residues were predicted to lie at the interface of at least one structure for WT/Delta (spike protein sites 403-406,409,414-417,419-421,445-447, 449,453,455-461,473-478,484-498,500-506), 56 for Gamma (WT/Delta+408,439,480,499), and 53 for Omicron (WT/Delta+439,499,-445). We introduced all of the 19 possible mutations at each interface site (all 56 identified for any variant) for these 100 structures for the WT, Delta, Gamma, and Omicron variants with the exception of WT reversion for the variants. Mutations were introduced using the PackRotamersMover with design specified by the input resfile (see Data Availability for example). Only the targeted residue was modified: side chain conformations for all other residues were fixed and no backbone minimization was applied before filtering for candidates of interest as described below.

Introducing the mutation in this way without repacking and minimization results in many unrealistic, high-energy conformations which are difficult or impossible to interpret without further optimization. The benefit is speed. This procedure can be executed in under 30 seconds per structure while global repacking and minimization may take hours. An alternative approach is to apply backbone minimization and sidechain repacking within a local region of the structure centered at the modified site as implemented within Rosetta through the Flex ddG protocol(44). This approach has been used to accurately predict mutations conferring increased infectivity for SARS-CoV-2(45) and rationally design NAb against the antigen(46, 47) but is significantly slower than introducing mutations without, even local, optimization. With or without local optimization prior to filtering, global repacking and minimization may be desired to confirm top candidates. While our approach may introduce a higher false-negative rate than what one would achieve applying local optimization prior to filtering, it maximizes the breadth of candidate mutations considered and is able to recapitulate experimental results.

The total score and dG separated of the mutant structures were computed. Unlike in earlier steps where the unbound state was repacked during the dG separated calculation, no repacking was conducted for the reasons described above. Each mutant structure was matched with its initial WT/variant conformation and the change in dG separated, *(S*_*M*_^*C*^*-S*_*M*_^*U*^*)-(S*_*WT*_^*C*^*-S*_*WT*_^*U*^), and total score, *S*_*M*_^*C*^*-S*_*WT*_^*C*^, were calculated. The average of each value taken over the ensemble of the 50 RBD-ACE2 conformations was compared with experimentally determined ACE2 binding affinity for single RBD mutants(12). While the change in dG separated is technically the ddG for the mutant and may be expected to correspond to binding affinity, we find *(S*_*M*_^*C*^*-S*_*M*_^*U*^*)-(S*_*WT*_ ^*C*^*-S*_*WT*_ ^*U*^) over this ensemble is not generally predictive of ACE2 affinity as *(S*_*M*_^*C*^*-S*_*M*_^*U*^*)-(S*_*WT*_ ^*C*^*-S*_*WT*_^*U*^) is approximately zero for many mutants (Figure S14). This may be expected without repacking and minimization. Nonetheless, when nonzero, the change in dG separated compares favorably with ACE2 affinity (see below).

The change in total score is less sensitive and an adequate predictor of affinity (Figure S15) within the range *S*_*M*_^*C*^*-S*_*WT*_ ^*C*^*<5* such that the minimum relative binding affinity decreases with increasing change in total score. Additional validation that *S*_*M*_^*C*^*-S*_*WT*_ ^*C*^ is a reasonable predictor of binding affinity, below some threshold, may be obtained through demonstrating *S*_*M*_^*C*^*-S*_*WT*_ ^*C*^ for each mutant is highly correlated between the RBD-ACE2 complex and the RBD-NAb complex. Furthermore, when the difference between the change in dG separated for the NAb complex and the dG separated for the ACE2 complex, *[(S*_*M*_^*C*^*-S*_*M*_^*U*^*)-(S*_*WT*_ ^*C*^*-S*_*WT*_^*U*^)]_*NAb*_*-[(S*_*M*_^*C*^*-S*_*M*_^*U*^*)-(S*_*WT*_ ^*C*^*-S*_*WT*_ ^*U*^)]_*ACE2*_, is positive, mutants lie above the identity line (y==x). The reverse is true for mutants falling below the identity line (Figures 1C/3B). In other words, on average, mutants in the RBD-ACE2 complex behave similarly to mutants in the RBD-NAb complex (which is what one would expect for realistic models of interfaces with an overlapping footprint). When differences do appear, they correspond to interactions with the RBD which are specific to the binding partner (ACE2/NAb) and are correctly reflected by the change in dG separated.

### Directly Assessing Additive Effects Vs Epistasis

In the main text, we demonstrate that most mutations appear in a similar position of the receptor cost / antibody cost phase space for the WT and Gamma/Delta variants. Consequently, the ensemble of vaccine escape candidates is largely conserved. Whether epistatic effects due to multiple mutations are present or if the impact of multiple mutations is simply additive can be more directly assessed for each complex by plotting the change in the total score after introducing each mutation in the variant against the change in total score after introducing each mutation in the WT. Figure S5 displays the change in total score for each interface mutation relative to the WT and all variants for both ACE2 and the NAb. The minimum value (less 1) is subtracted from each distribution. The effect of each mutation at the interface in the WT is highly correlated for the Delta variant. This correlation is observed for the Gamma and Omicron variants as well with more significant variation as expected. There is additionally a small systematic bias towards greater NAb destabilization for mutations with small changes in the total score for the Omicron variant. As discussed in the main text, while this epistatic signature is not observed for the Gamma variant, this trend did not significantly alter the landscape of escape-exacerbating mutations.

### Alternative Construction of Variant Ensembles

Beginning with the conformational ensemble constructed for the WT, we introduced the variant mutations 452R|478K for the Delta variant, 417T|484K|501Y for the Gamma variant, and 339D|371L|373P|375F|417N|440K|446S|477N|478K|484A|493R|496S| 498R|501Y|505H for the Omicron variant and completed 5 rounds of the FastRelaxMover (15 for the Omicron variant) using default parameters (with the exception of disabling design) as described above. The resulting values for the total score and dG separated were comparable to that of the ensembles constructed for the variants beginning with the WT crystal structure (Figures S16-18); however, some structures displayed footprints more similar to the WT than those of the reference ensemble constructed as described in the main text. Thus, this computationally cheap alternative may be utilized for rapid evaluation; but the more comprehensive exploration of the conformational space described in the main is likely desired in many circumstances.

## Data Availability

The data has been deposited through Zenodo(48) (https://doi.org/10.5281/zenodo.5297698) including GISAID acknowledgements. Previously published data were used for this work: GISAID(23). Data is additionally made available through FTP: https://ftp.ncbi.nih.gov/pub/wolf/_suppl/SARSstruct21/

## Author contributions

NDR, and GF collected data; NDR, YIW, GF, PF, FZ, and EVK analyzed data; NDR and EVK wrote the manuscript that was edited and approved by all authors.

## Acknowledgements

The authors thank the members of Jeff Gray’s research group from Johns Hopkins University, in particular, Rahel Frick, for their consultation regarding best practices using the Rosetta modelling software. The authors also thank Koonin group members for helpful discussions. NDR, YIW, and EVK are supported by the Intramural Research Program of the National Institutes of Health (National Library of Medicine).

## References

1. Zhan SH, Deverman BE, Chan YA. SARS-CoV-2 is well adapted for humans. What does this mean for re-emergence? BioRxiv. 2020.

2. van Dorp L, Richard D, Tan CC, Shaw LP, Acman M, Balloux F. No evidence for increased transmissibility from recurrent mutations in SARS-CoV-2. Nature communications. 2020;11(1):1–8.

3. Rochman ND, Wolf YI, Faure G, Mutz P, Zhang F, Koonin EV. Ongoing global and regional adaptive evolution of SARS-CoV-2. Proceedings of the National Academy of Sciences. 2021;118(29).

4. Rochman N, Wolf Y, Koonin EV. Evolution of human respiratory virus epidemics. F1000Research. 2021;10.

5. Saad-Roy CM, Wagner CE, Baker RE, Morris SE, Farrar J, Graham AL, et al. Immune life history, vaccination, and the dynamics of SARS-CoV-2 over the next 5 years. Science. 2020;370(6518):811–8.

6. Amanat F, Krammer F. SARS-CoV-2 vaccines: status report. Immunity. 2020;52(4):583–9.

7. Rochman N, Wolf Y, Koonin EV. Substantial impact of post-vaccination contacts on cumulative infections during viral epidemics. F1000Research. 2021;10.

8. Greaney AJ, Starr TN, Gilchuk P, Zost SJ, Binshtein E, Loes AN, et al. Complete mapping of mutations to the SARS-CoV-2 spike receptor-binding domain that escape antibody recognition. Cell host & microbe. 2021;29(1):44-57. e9.

9. Greaney AJ, Starr TN, Barnes CO, Weisblum Y, Schmidt F, Caskey M, et al. Mapping mutations to the SARS-CoV-2 RBD that escape binding by different classes of antibodies. Nature Communications. 2021;12(1):1–14.

10. Nelson G, Buzko O, Spilman PR, Niazi K, Rabizadeh S, Soon-Shiong PR. Molecular dynamic simulation reveals E484K mutation enhances spike RBD-ACE2 affinity and the combination of E484K, K417N and N501Y mutations (501Y. V2 variant) induces conformational change greater than N501Y mutant alone, potentially resulting in an escape mutant. BioRxiv. 2021.

11. Laffeber C, de Koning K, Kanaar R, Lebbink JH. Experimental evidence for enhanced receptor binding by rapidly spreading SARS-CoV-2 variants. Journal of Molecular Biology. 2021;433(15):167058.

12. Starr TN, Greaney AJ, Hilton SK, Ellis D, Crawford KH, Dingens AS, et al. Deep mutational scanning of SARS-CoV-2 receptor binding domain reveals constraints on folding and ACE2 binding. Cell. 2020;182(5):1295-310. e20.

13. Leaver-Fay A, Tyka M, Lewis SM, Lange OF, Thompson J, Jacak R, et al. ROSETTA3: an object-oriented software suite for the simulation and design of macromolecules. Methods in enzymology. 2011;487:545–74.

14. Li B, Deng A, Li K, Hu Y, Li Z, Xiong Q, et al. Viral infection and transmission in a large well-traced outbreak caused by the Delta SARS-CoV-2 variant. MedRxiv. 2021.

15. Duong D. Alpha, Beta, Delta, Gamma: What’s important to know about SARS-CoV-2 variants of concern? : Can Med Assoc; 2021.

16. Lan J, Ge J, Yu J, Shan S, Zhou H, Fan S, et al. Structure of the SARS-CoV-2 spike receptor-binding domain bound to the ACE2 receptor. Nature. 2020;581(7807):215–20.

17. Jiang S, Hillyer C, Du L. Neutralizing antibodies against SARS-CoV-2 and other human coronaviruses. Trends in immunology. 2020;41(5):355–9.

18. Alford RF, Leaver-Fay A, Jeliazkov JR, O’Meara MJ, DiMaio FP, Park H, et al. The Rosetta all-atom energy function for macromolecular modeling and design. Journal of chemical theory and computation. 2017;13(6):3031–48.

19. Starr TN, Greaney AJ, Addetia A, Hannon WW, Choudhary MC, Dingens AS, et al. Prospective mapping of viral mutations that escape antibodies used to treat COVID-19. Science. 2021;371(6531):850–4.

20. Starr TN, Greaney AJ, Dingens AS, Bloom JD. Complete map of SARS-CoV-2 RBD mutations that escape the monoclonal antibody LY-CoV555 and its cocktail with LY-CoV016. Cell Reports Medicine. 2021;2(4):100255.

21. Wang Z, Schmidt F, Weisblum Y, Muecksch F, Barnes CO, Finkin S, et al. mRNA vaccine-elicited antibodies to SARS-CoV-2 and circulating variants. Nature. 2021;592(7855):616–22.

22. Greaney AJ, Loes AN, Crawford KH, Starr TN, Malone KD, Chu HY, et al. Comprehensive mapping of mutations in the SARS-CoV-2 receptor-binding domain that affect recognition by polyclonal human plasma antibodies. Cell host & microbe. 2021;29(3):463-76. e6.

23. Shu Y, McCauley J. GISAID: Global initiative on sharing all influenza data–from vision to reality. Eurosurveillance. 2017;22(13):30494.

24. Zhang Q, Ju B, Ge J, Chan JF-W, Cheng L, Wang R, et al. Potent and protective IGHV3-53/3-66 public antibodies and their shared escape mutant on the spike of SARS-CoV-2. Nature communications. 2021;12(1):1–12.

25. Alenquer M, Ferreira F, Lousa D, Valerio M, Medina-Lopes M, Bergman M-L, et al. Amino acids 484 and 494 of SARS-CoV-2 spike are hotspots of immune evasion affecting antibody but not ACE2 binding. bioRxiv. 2021.

26. Breen MS, Kemena C, Vlasov PK, Notredame C, Kondrashov FA. Epistasis as the primary factor in molecular evolution. Nature. 2012;490(7421):535–8.

27. Sakai W, Swisher EM, Karlan BY, Agarwal MK, Higgins J, Friedman C, et al. Secondary mutations as a mechanism of cisplatin resistance in BRCA2-mutated cancers. Nature. 2008;451(7182):1116–20.

28. Levin BR, Perrot V, Walker N. Compensatory mutations, antibiotic resistance and the population genetics of adaptive evolution in bacteria. Genetics. 2000;154(3):985–97.

29. Rochman ND, Wolf YI, Koonin EV. Deep phylogeny of cancer drivers and compensatory mutations. Communications biology. 2020;3(1):1–11.

30. Kupferschmidt K. Where did ‘weird’Omicron come from? : American Association for the Advancement of Science; 2021.

31. Wilhelm A, Widera M, Grikscheit K, Toptan T, Schenk B, Pallas C, et al. Reduced Neutralization of SARS-CoV-2 Omicron Variant by Vaccine Sera and monoclonal antibodies. medRxiv. 2021.

32. Gruell H, Vanshylla K, Tober-Lau P, Hillus D, Schommers P, Lehmann C, et al. mRNA booster immunization elicits potent neutralizing serum activity against the SARS-CoV-2 Omicron variant. medRxiv. 2021.

33. Zhang Y, Zhang J, Chen Y, Luo B, Yuan Y, Huang F, et al. The ORF8 protein of SARS-CoV-2 mediates immune evasion through potently downregulating MHC-I. BioRxiv. 2020.

34. Zinzula L. Lost in deletion: The enigmatic ORF8 protein of SARS-CoV-2. Biochemical and biophysical research communications. 2021;538:116–24.

35. Garushyants SK, Rogozin IB, Koonin EV. Template switching and duplications in SARS-CoV-2 genomes give rise to insertion variants that merit monitoring. Communications Biology. 2021;4(1):1–9.

36. Zhang J, Cai Y, Xiao T, Lu J, Peng H, Sterling SM, et al. Structural impact on SARS-CoV-2 spike protein by D614G substitution. Science. 2021;372(6541):525–30.

37. Hurlburt NK, Seydoux E, Wan Y-H, Edara VV, Stuart AB, Feng J, et al. Structural basis for potent neutralization of SARS-CoV-2 and role of antibody affinity maturation. Nature communications. 2020;11(1):1–7.

38. Seydoux E, Homad LJ, MacCamy AJ, Parks KR, Hurlburt NK, Jennewein MF, et al. Analysis of a SARS-CoV-2-infected individual reveals development of potent neutralizing antibodies with limited somatic mutation. Immunity. 2020;53(1):98-105. e5.

39. Voss WN, Hou YJ, Johnson NV, Delidakis G, Kim JE, Javanmardi K, et al. Prevalent, protective, and convergent IgG recognition of SARS-CoV-2 non-RBD spike epitopes. Science. 2021;372(6546):1108–12.

40. Tyka MD, Keedy DA, André I, DiMaio F, Song Y, Richardson DC, et al. Alternate states of proteins revealed by detailed energy landscape mapping. Journal of molecular biology. 2011;405(2):607–18.

41. Fleishman SJ, Leaver-Fay A, Corn JE, Strauch E-M, Khare SD, Koga N, et al. RosettaScripts: a scripting language interface to the Rosetta macromolecular modeling suite. PloS one. 2011;6(6):e20161.

42. Tovchigrechko A, Vakser IA. How common is the funnel-like energy landscape in protein-protein interactions? Protein science. 2001;10(8):1572–83.

43. Grünberg R, Nilges M, Leckner J. Flexibility and conformational entropy in protein-protein binding. Structure. 2006;14(4):683–93.

44. Barlow KA, ÓConchúir S, Thompson S, Suresh P, Lucas JE, Heinonen M, et al. Flex ddG: Rosetta ensemble-based estimation of changes in protein–protein binding affinity upon mutation. The Journal of Physical Chemistry B. 2018;122(21):5389–99.

45. Xue T, Wu W, Guo N, Wu C, Huang J, Lai L, et al. Single point mutations can potentially enhance infectivity of SARS-CoV-2 revealed by in silico affinity maturation and SPR assay. RSC Advances. 2021;11(24):14737–45.

46. Riahi S, Lee JH, Wei S, Cost R, Masiero A, Prades C, et al. Application of an integrated computational antibody engineering platform to design SARS-CoV-2 neutralizers. bioRxiv. 2021.

47. Desautels T, Zemla A, Lau E, Franco M, Faissol D. Rapid in silico design of antibodies targeting SARS-CoV-2 using machine learning and supercomputing. BioRxiv. 2020.

48. Epistasis at the SARS-CoV-2 RBD Interface and the Propitiously Boring Implications for Vaccine Escape [Data set] [Internet]. 2021.

