## Supplementary figures, tables and code for "Epistasis at the SARS-CoV-2 RBD Interface and the Propitiously Boring Implications for Vaccine Escape"

**This PDF file includes:**

Figures S1 to S18
Table S1

**Additional supplementary information can be found:**

https://ftp.ncbi.nih.gov/pub/wolf/_suppl/SARSevo21/
**or**
https://doi.org/10.5281/zenodo.5297698

This repository includes:

GISAID Acknowledgements
An example resfile
RosettaScripts .xml files
Complex conformations (50 each) for WT/Gamma/Delta; antibody (NAb) and receptor (ACE2)

For correspondence:

Nash Rochman

Feng Zhang

Eugene Koonin

### Supplemental Figures

**Figure S1.** Number of escape candidates and the mean score for each candidate, *max*($[S_M^C - S_{WT}^C]_{NAb} - [S_M^C - S_{WT}^C]_{ACE2}$, $\Delta\Delta G_{NAb} - \Delta\Delta G_{ACE2}$), across all RBD Interface sites for the WT.

**Figure S2.** Classical multidimensional scaling (CMDS) applied to the pairwise interface RMSDs among all RBD-ACE2 and RBD-NAb complexes (over spike protein sites 403-406,408-409,414-417,419-421,439,445-447,449,453,455-461,473-478,480,484-506).

**Figure S3.** *Left:* Receptor cost for mutations at the interface that were experimentally demonstrated to reduce NAb activity in the WT(1) (right) and other experimentally studied mutations at the interface (left). P-values for a Wilcoxon Rank Sum Test between the left WT distribution and all others shown. Right: Distribution of median values over 1000x bootstrap. P-values between left and right distributions for the WT and all variants shown.

**Figure S4.** Movement within the plane of receptor cost vs. antibody cost. Each arrow represents an amino acid substitution in site, **A** 408, **B** 504, or **C** 446 and arrows point from the position on the plane corresponding to substitution in the WT to the position corresponding to the substitution in each variant. Color represents the sign of $\Delta\Delta G_{NAb} - \Delta\Delta G_{ACE2}$ (blue, negative; red, positive).

**Figure S5.** Total score change induced by mutation in each variant vs the WT. The minimum value (less 1) is subtracted from each distribution.

**Figure S6.** dG separated vs. the interface RMSD with the structure corresponding to the lowest dG separated for the WT computed over the alpha carbon backbone atoms of identified interface residues and the associated (+/-) 3 amino acid neighborhoods of each in the RBD (spike protein sites 400-424, 440-464, and 470-509).

**Figure S7.** dG separated vs. the interface RMSD with the structure corresponding to the lowest dG separated for the Delta variant computed over the alpha carbon backbone atoms of identified interface residues and the associated (+/-) 3 amino acid neighborhoods of each in the RBD (spike protein sites 400-424, 436-464, and 470-509).

**Figure S8.** dG separated vs. the interface RMSD with the structure corresponding to the lowest dG separated for the Gamma variant computed over the alpha carbon backbone atoms of identified interface residues and the associated (+/-) 3 amino acid neighborhoods of each in the RBD (spike protein sites 400-424, 434-464, and 470-509).

**Figure S9.** dG separated vs. the interface RMSD with the structure corresponding to the lowest dG separated for the Omicron variant computed over the alpha carbon backbone atoms of identified interface residues and the associated (+/-) 3 amino acid neighborhoods of each in the RBD (spike protein sites 400-424, 434-464, and 470-510).

**Figure S10.** dG separated vs. the total score for WT conformations achieved after iterative applications of FastRelax beginning with the crystal structure. Blue points represent iterations 1-4 and green represent iterations 5-12. Points outlined in black are the 50 conformations selected to represent each complex.

**Figure S11.** dG separated vs. the total score for Delta variant conformations achieved after iterative applications of FastRelax beginning with the crystal structure. Blue points represent iterations 1-4 and green represent iterations 5-12. Points outlined in black are the 50 conformations selected to represent each complex. Gray dots are WT conformations.

**Figure S12.** dG separated vs. the total score for Gamma variant conformations achieved after iterative applications of FastRelax beginning with the crystal structure. Blue points represent iterations 1-4 and green represent iterations 5-12. Points outlined in black are the 50 conformations selected to represent each complex. Gray dots are WT conformations.

**Figure S13.** dG separated vs. the total score for Omicron variant conformations achieved after iterative applications of FastRelax beginning with the crystal structure. Blue points represent iterations 1-4 and green represent iterations 5-15. Points outlined in black are the 50 conformations selected to represent each complex. Gray dots are WT conformations.

**Figure S14.** Relative ACE2 affinity as reported by Starr et. al(2), positive values indicate stronger affinity, vs difference in dG separated between the mutant and the WT.

**Figure S15.** Relative ACE2 affinity as reported by Starr et. al(2), positive values indicate stronger affinity, vs difference in total score between the mutant and the WT.

**Figure S16.** dG separated vs total score for the original Delta variant conformational ensemble generated and the alternate ensemble.

**Figure S17.** dG separated vs total score for the original Gamma variant conformational ensemble generated and the alternate ensemble.

**Figure S18.** dG separated vs total score for the original Omicron variant conformational ensemble generated and the alternate ensemble.

#### Supplemental Tables

**Table S1.** RosettaScripts files, descriptions, and example command line implementations.

#### References

1. Greaney AJ, Starr TN, Gilchuk P, Zost SJ, Binshtein E, Loes AN, et al. Complete mapping of mutations to the SARS-CoV-2 spike receptor-binding domain that escape antibody recognition. Cell host & microbe. 2021;29(1):44-57. e9.
2. Starr TN, Greaney AJ, Hilton SK, Ellis D, Crawford KH, Dingens AS, et al. Deep mutational scanning of SARS-CoV-2 receptor binding domain reveals constraints on folding and ACE2 binding. Cell. 2020;182(5):1295-310. e20.

**S1**

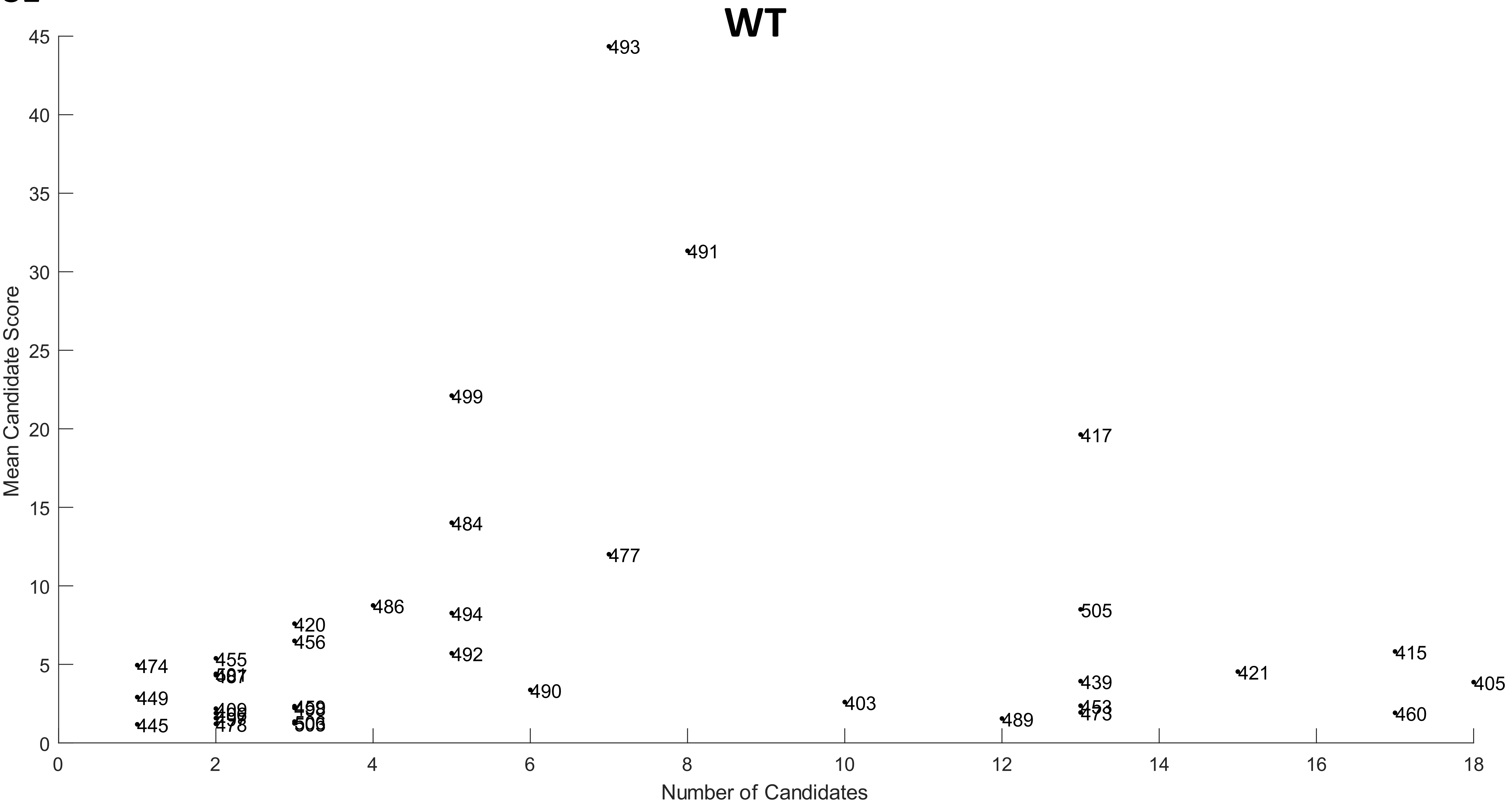
WT
Mean Candidate Score
Number of Candidates
0
5
10
15
20
25
30
35
40
45
0
2
4
6
8
10
12
14
16
18
493
491
499
417
484
477
486
505
420
494
456
492
415
474
455
421
439
405
490
449
403
489
460
445
478
473

S2

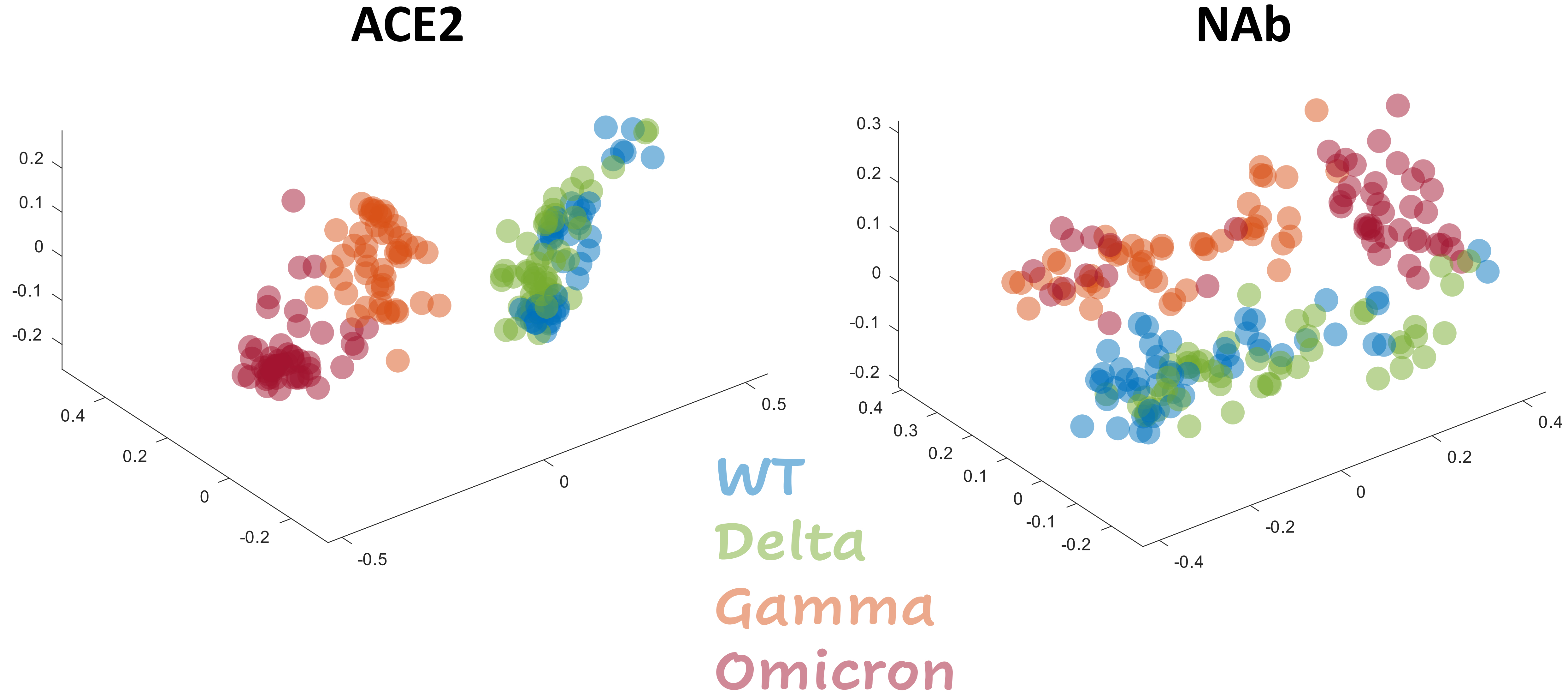
ACE2
NAb
0.2
0.1
0
-0.1
-0.2
0.4
0.2
0
-0.2
-0.5
0
0.5
0.3
0.2
0.1
0
-0.1
-0.2
0.4
0.3
0.2
0.1
0
-0.1
-0.2
-0.4
-0.2
0
0.2
0.4
WT
Delta
Gamma
Omicron

**S3**

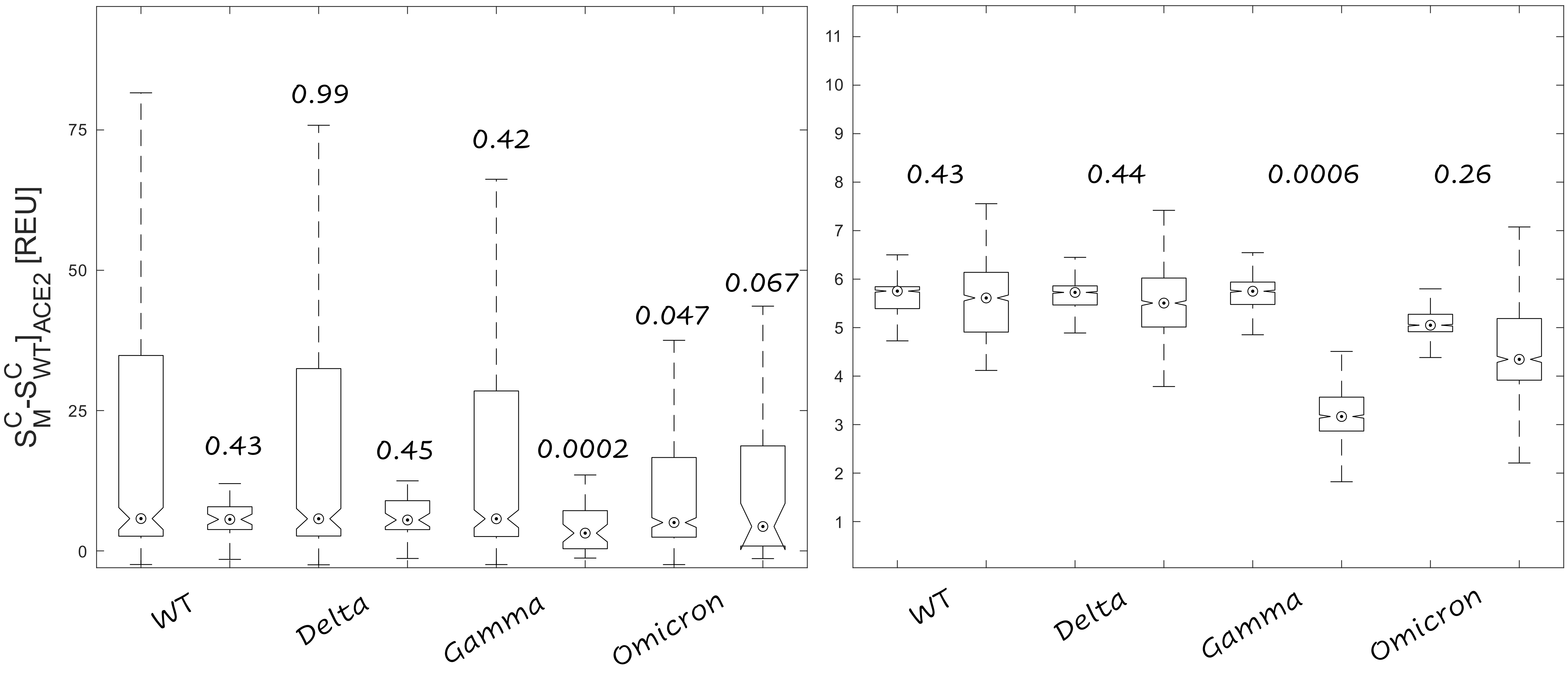

$S^C_M - S^C_{WT}]_{ACE2}$ [REU]
75
50
25
0
0.99
0.42
0.43
0.45
0.0002
0.047
0.067
WT
Delta
Gamma
Omicron
11
10
9
8
7
6
5
4
3
2
1
0.43
0.44
0.0006
0.26
WT
Delta
Gamma
Omicron

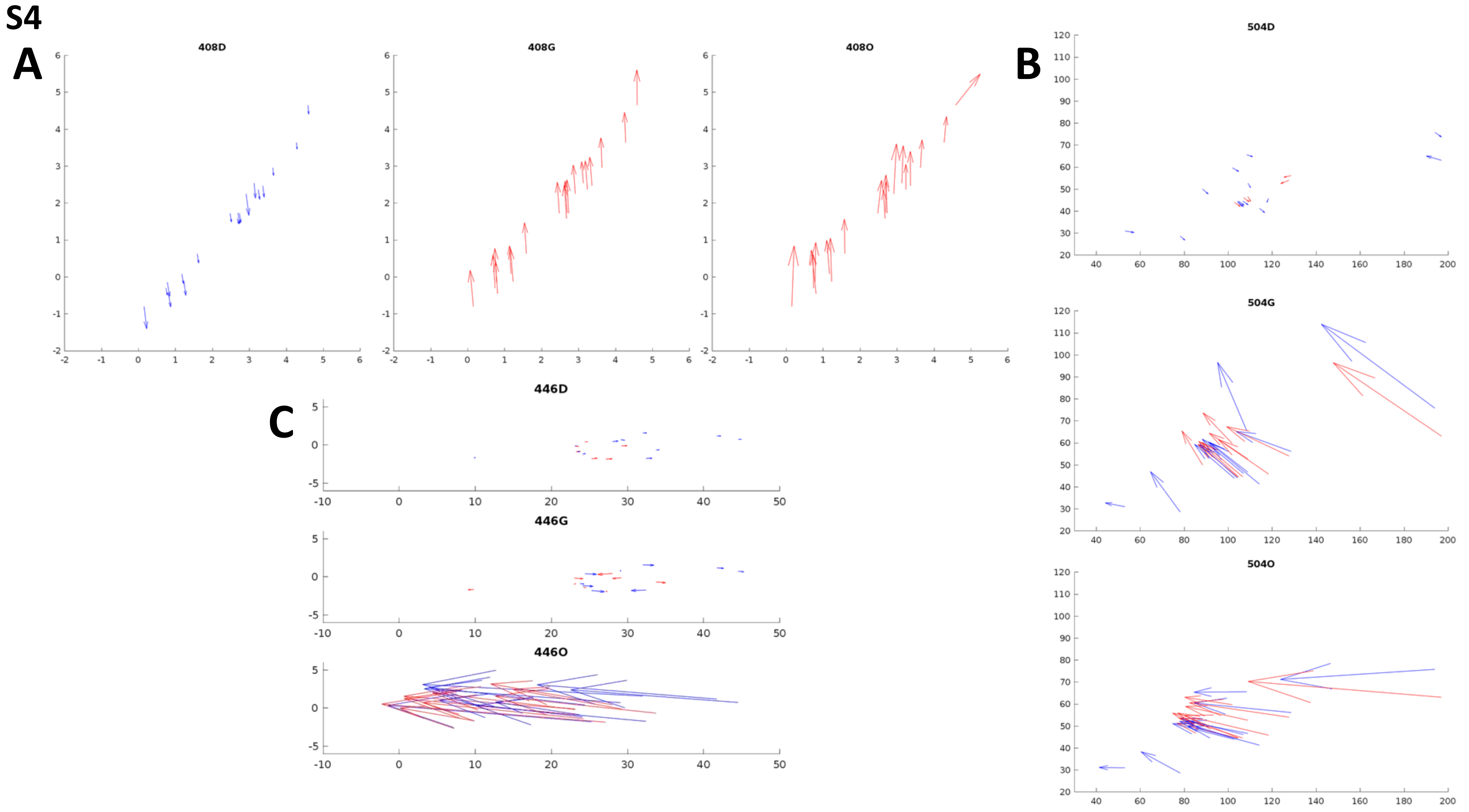

S4
A
408D
408G
408O
B
504D
504G
504O
C
446D
446G
446O

# S5

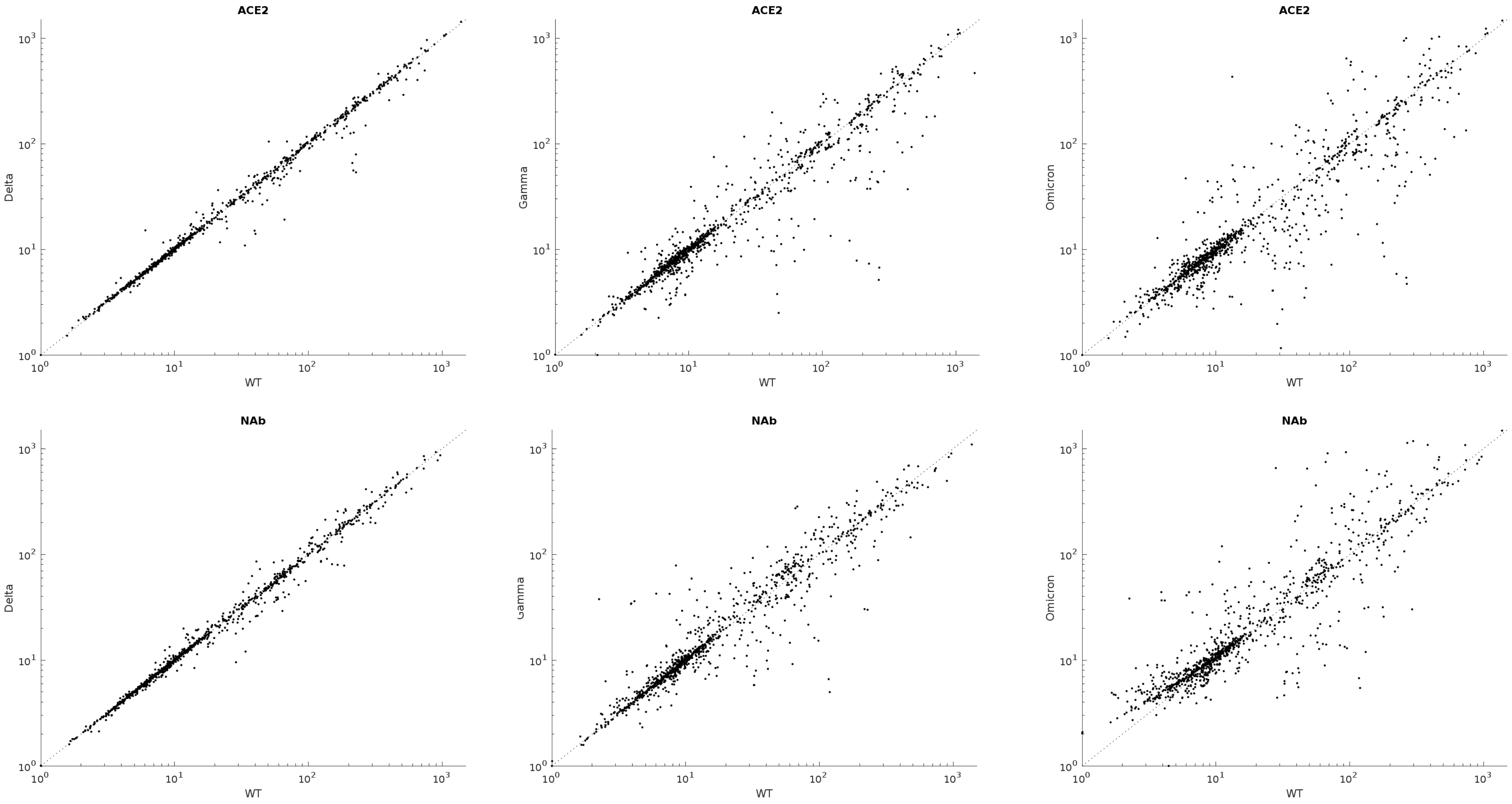
ACE2
Delta
WT
ACE2
Gamma
WT
ACE2
Omicron
WT
NAb
Delta
WT
NAb
Gamma
WT
NAb
Omicron
WT

**S6**

# WT

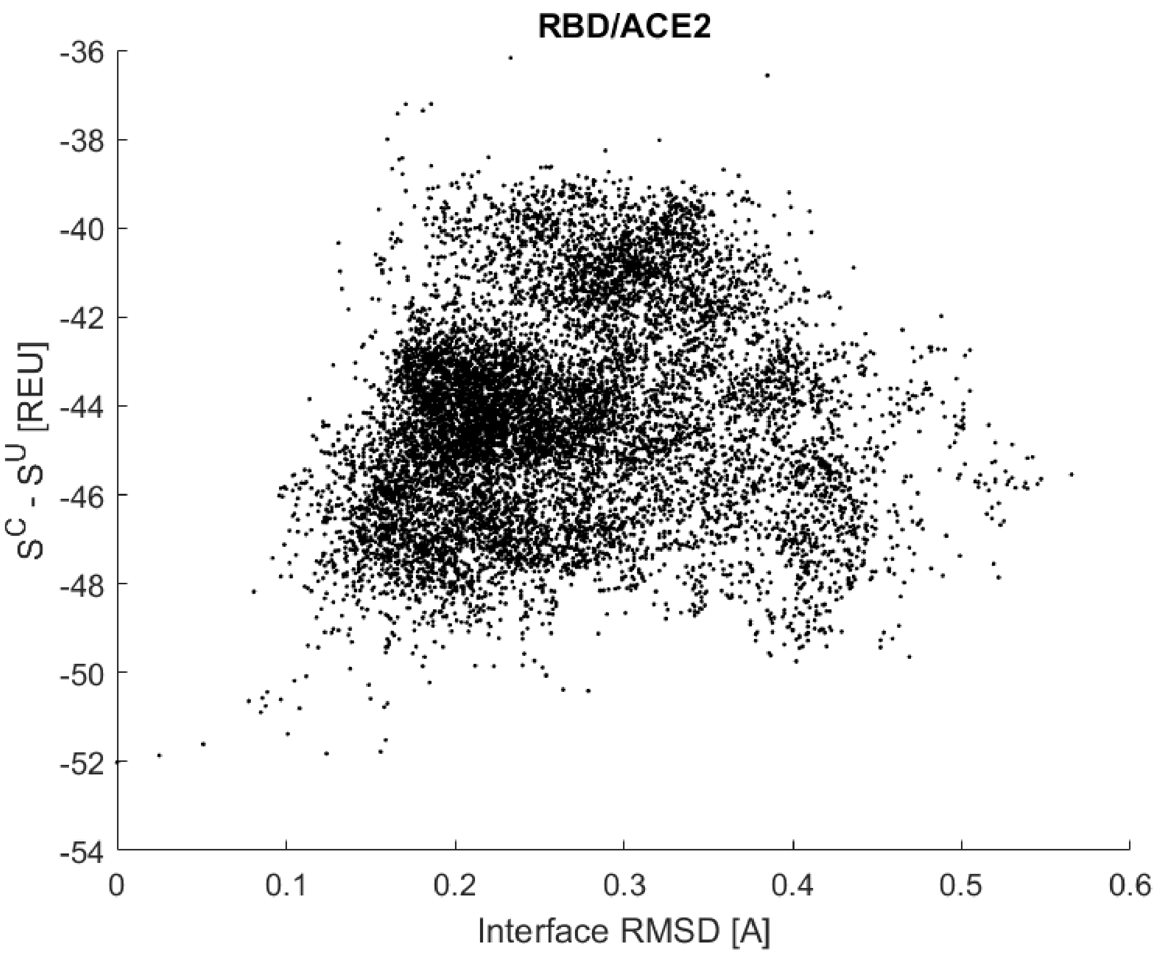

RBD/ACE2
-36
-38
-40
-42
-44
-46
-48
-50
-52
-54
$S^C - S^U$ [REU]
0
0.1
0.2
0.3
0.4
0.5
0.6
Interface RMSD [A]


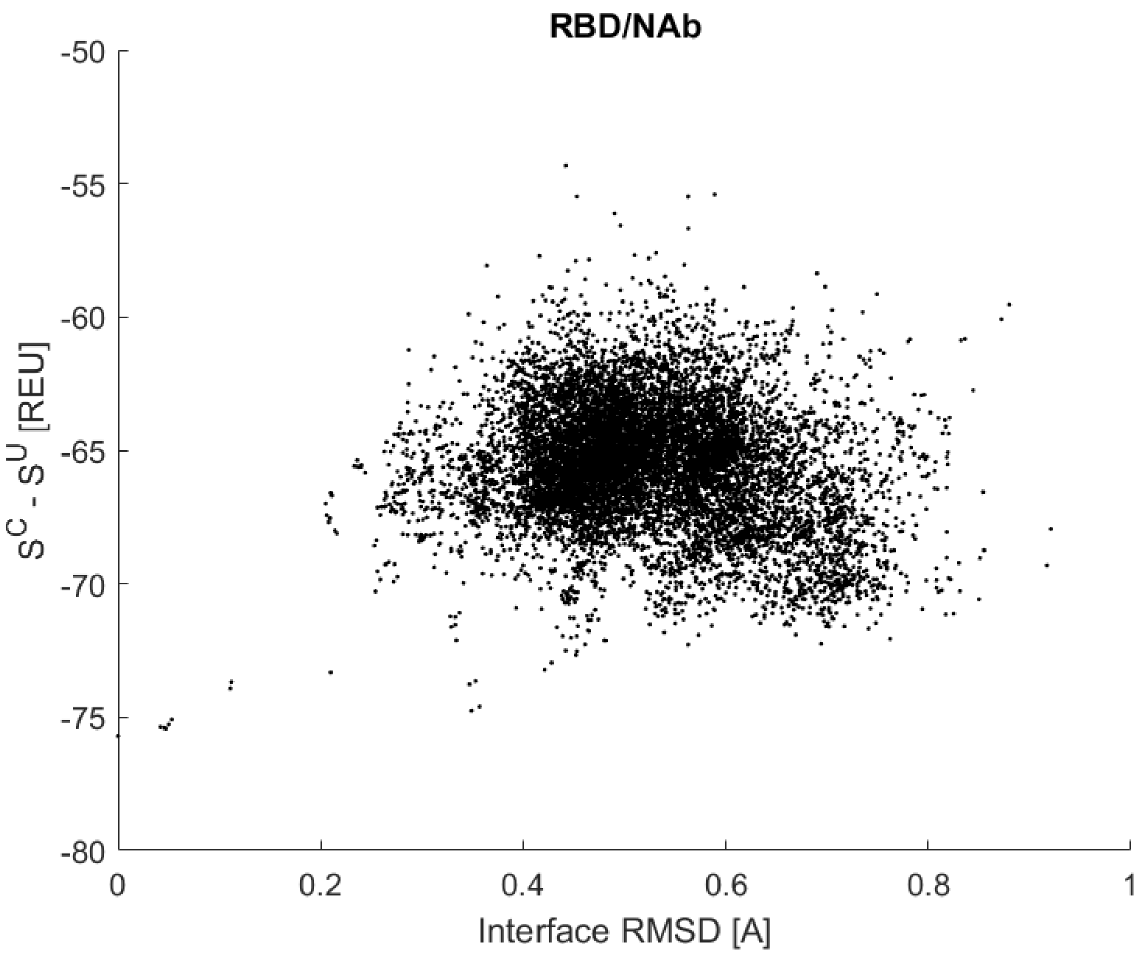

RBD/NAb
-50
-55
-60
-65
-70
-75
-80
$S^C - S^U$ [REU]
0
0.2
0.4
0.6
0.8
1
Interface RMSD [A]

# S7

#### Delta

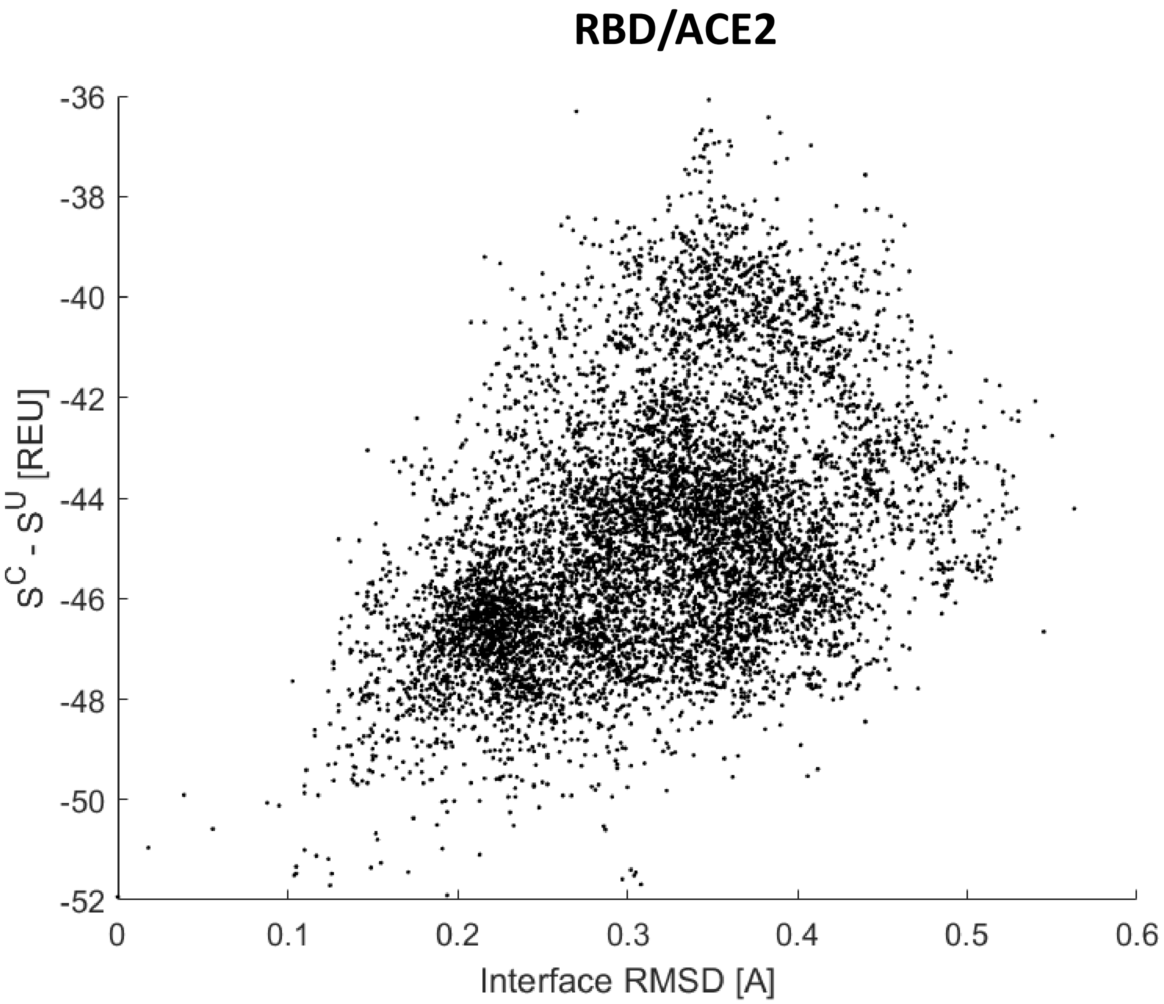
RBD/ACE2
-36
-38
-40
-42
-44
-46
-48
-50
-52
S^C - S^U [REU]
0
0.1
0.2
0.3
0.4
0.5
0.6
Interface RMSD [A]

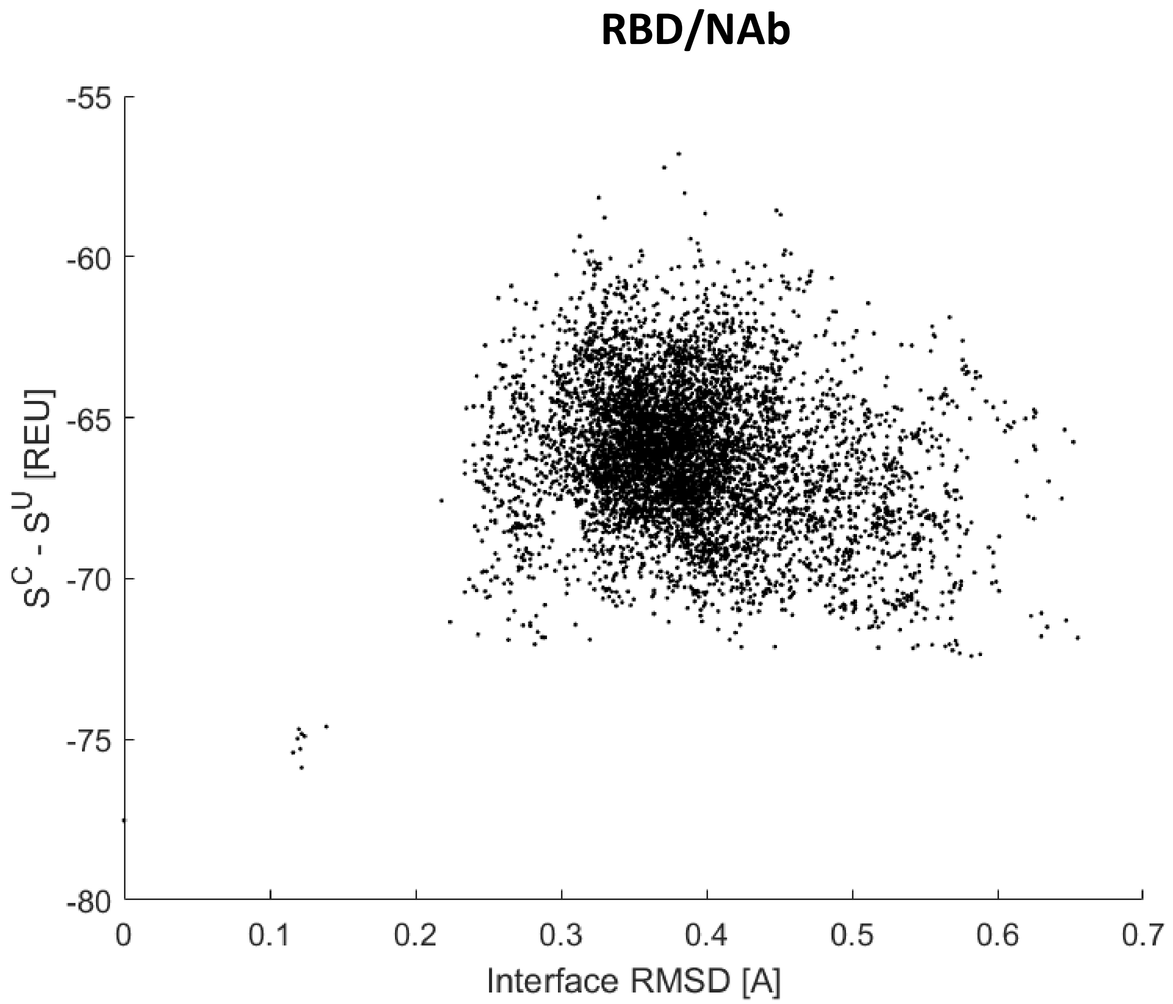
RBD/NAb
-55
-60
-65
-70
-75
-80
S^C - S^U [REU]
0
0.1
0.2
0.3
0.4
0.5
0.6
0.7
Interface RMSD [A]

**S8**

### Gamma

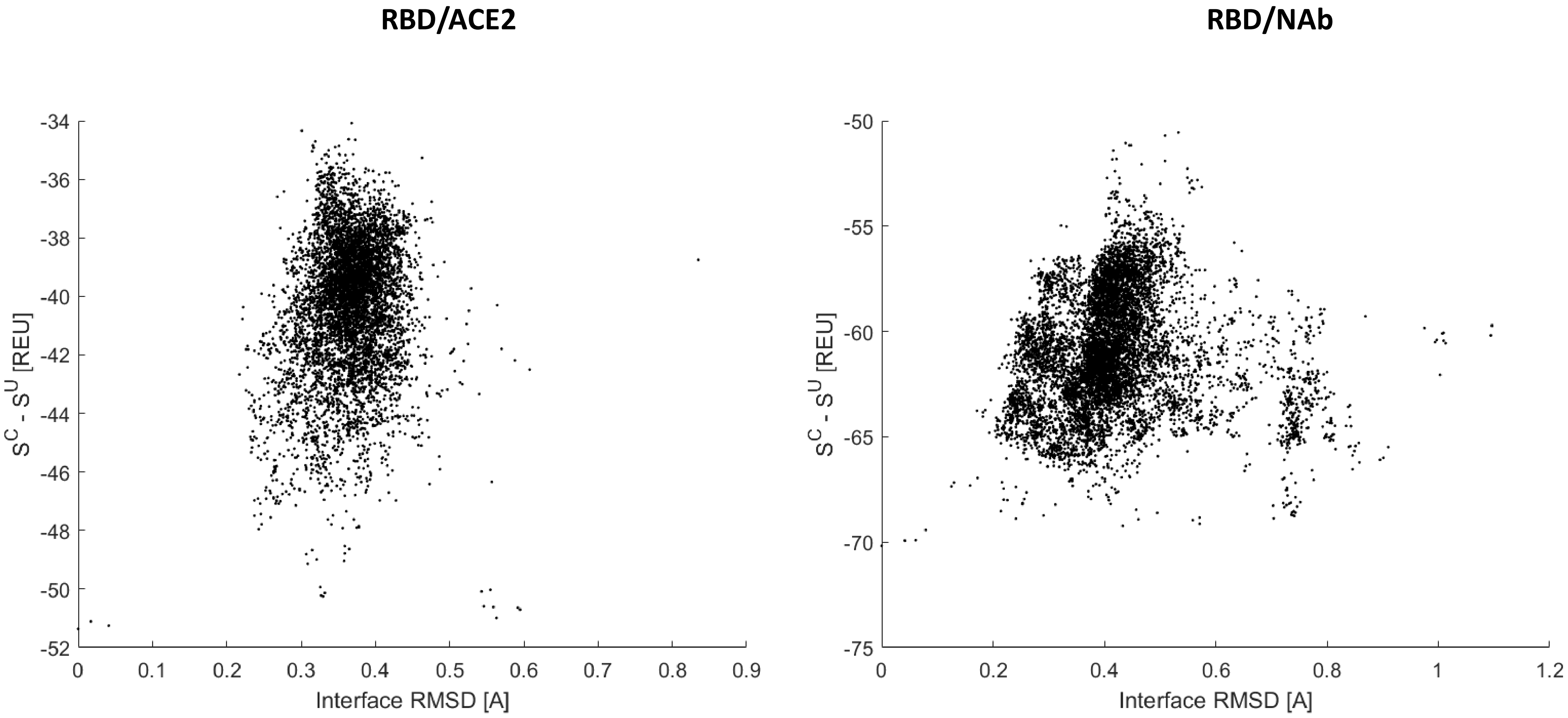
RBD/ACE2
RBD/NAb
$S^C - S^U$ [REU]
Interface RMSD [A]
-34
-36
-38
-40
-42
-44
-46
-48
-50
-52
0
0.1
0.2
0.3
0.4
0.5
0.6
0.7
0.8
0.9
-50
-55
-60
-65
-70
-75
0
0.2
0.4
0.6
0.8
1
1.2

**S9**

### Omicron

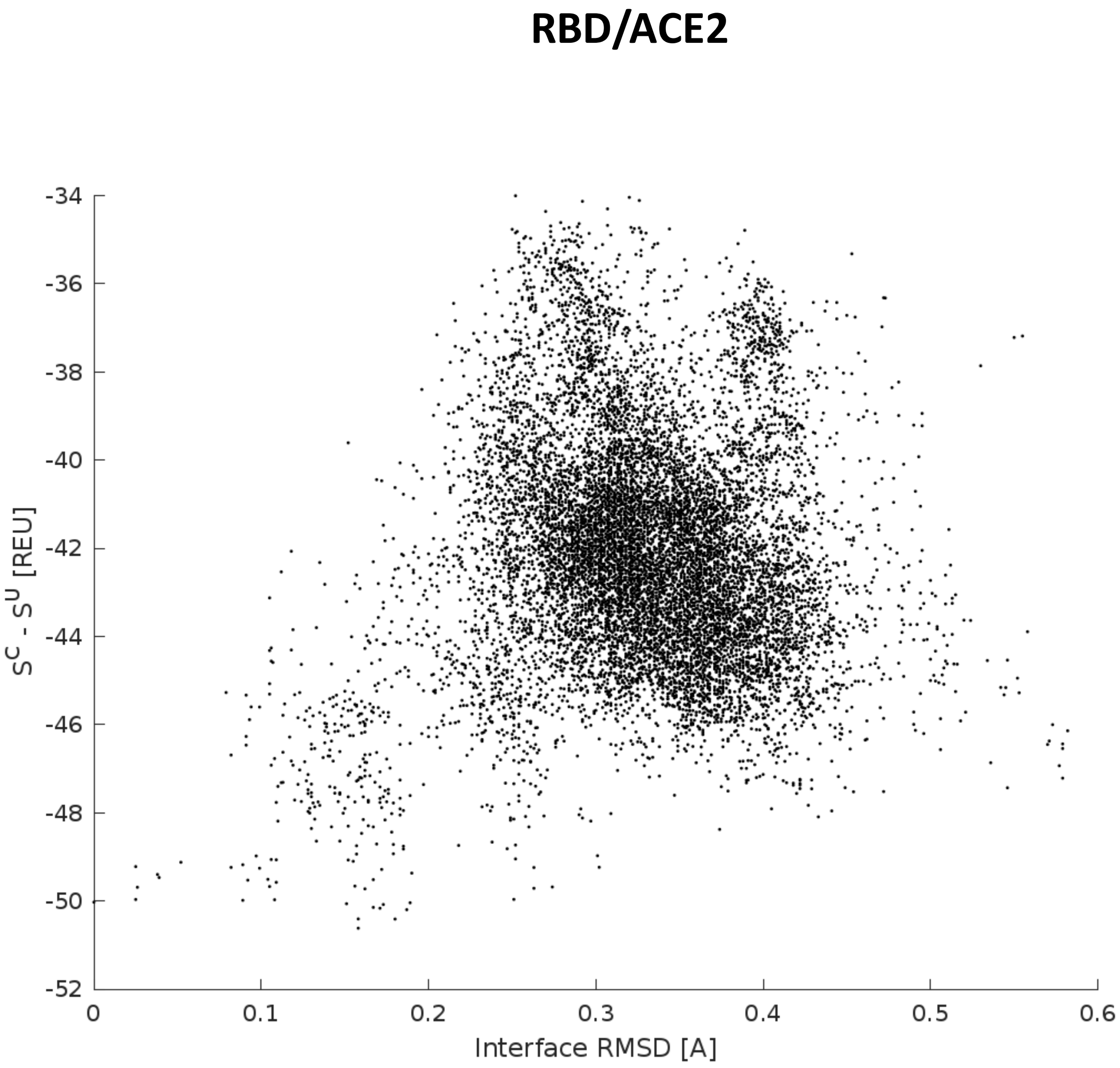
RBD/ACE2
-34
-36
-38
-40
-42
-44
-46
-48
-50
-52
$S^C - S^U$ [REU]
0
0.1
0.2
0.3
0.4
0.5
0.6
Interface RMSD [A]

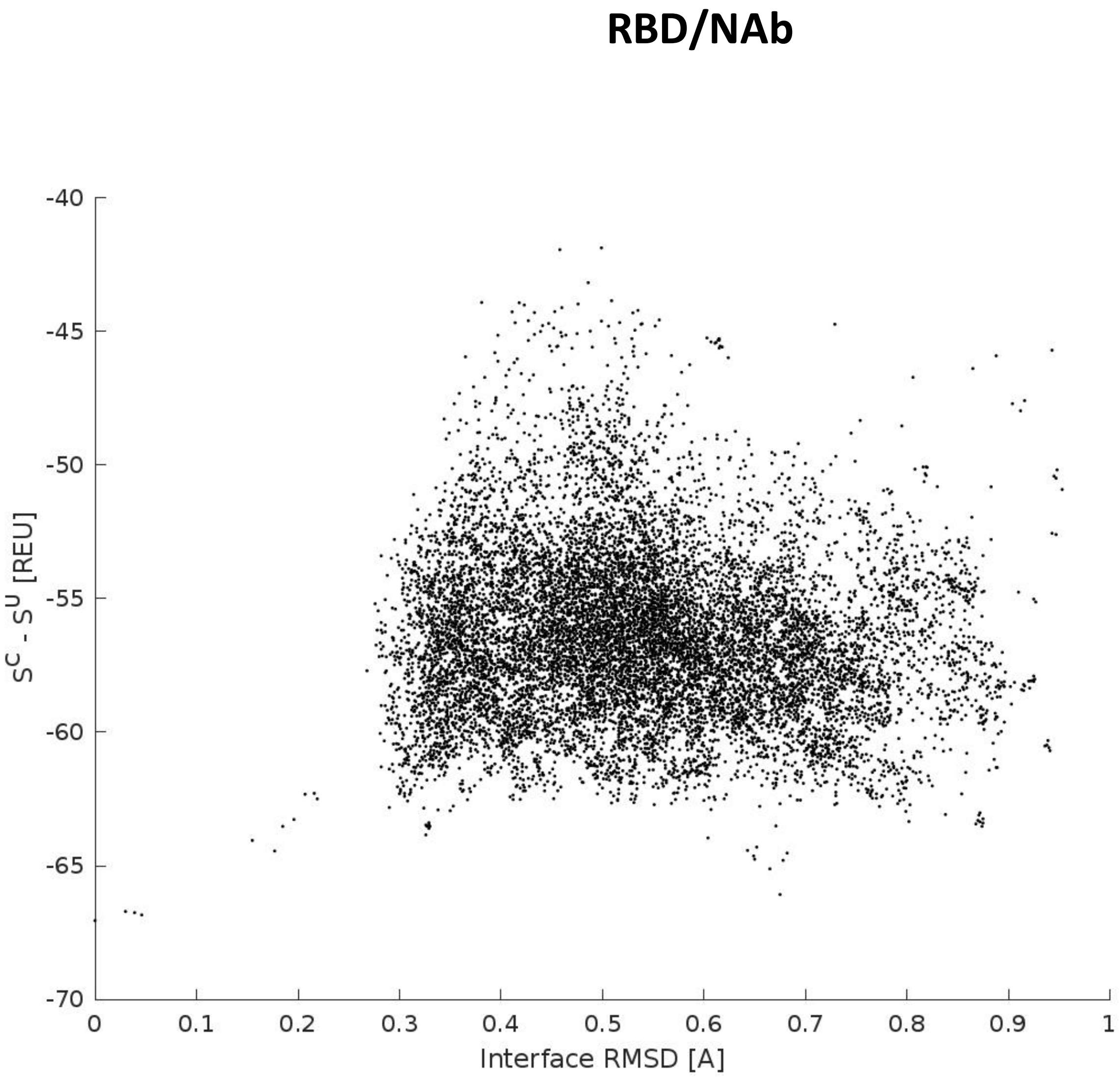
RBD/NAb
-40
-45
-50
-55
-60
-65
-70
$S^C - S^U$ [REU]
0
0.1
0.2
0.3
0.4
0.5
0.6
0.7
0.8
0.9
1
Interface RMSD [A]

**S10**

# WT

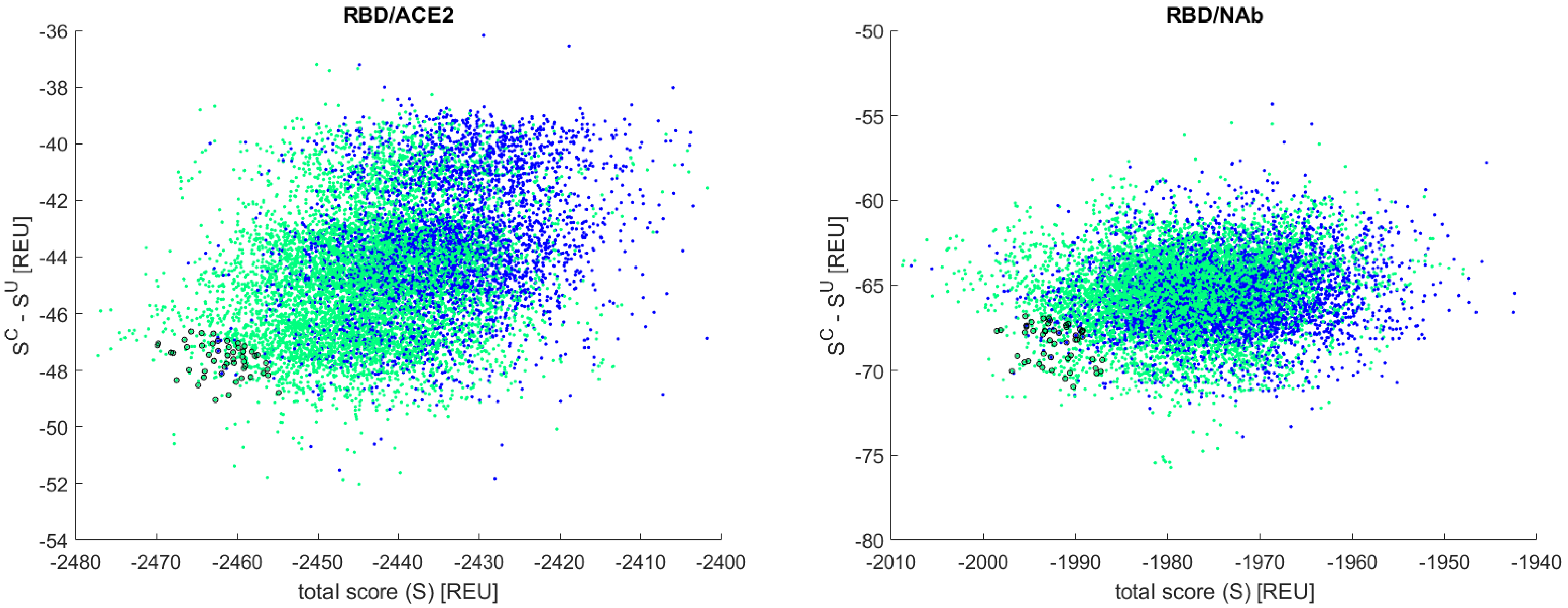
RBD/ACE2
-36
-38
-40
-42
-44
-46
-48
-50
-52
-54
$S^C - S^U$ [REU]
-2480
-2470
-2460
-2450
-2440
-2430
-2420
-2410
-2400
total score (S) [REU]
RBD/NAb
-50
-55
-60
-65
-70
-75
-80
$S^C - S^U$ [REU]
-2010
-2000
-1990
-1980
-1970
-1960
-1950
-1940
total score (S) [REU]

**S11**

### Delta

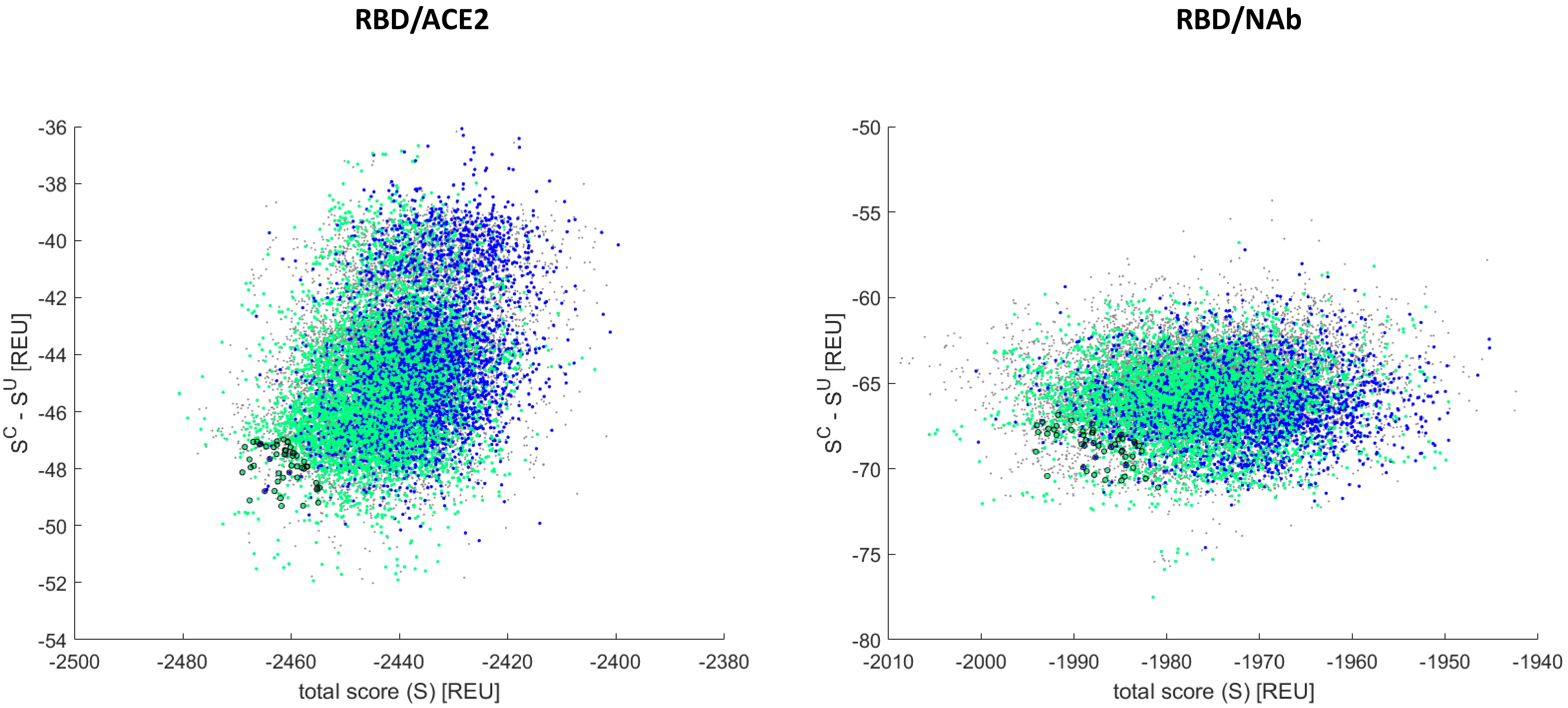
RBD/ACE2
RBD/NAb
$S^C - S^U$ [REU]
total score (S) [REU]
-36
-38
-40
-42
-44
-46
-48
-50
-52
-54
-2500
-2480
-2460
-2440
-2420
-2400
-2380
-50
-55
-60
-65
-70
-75
-80
-2010
-2000
-1990
-1980
-1970
-1960
-1950
-1940

**S12**

### Gamma

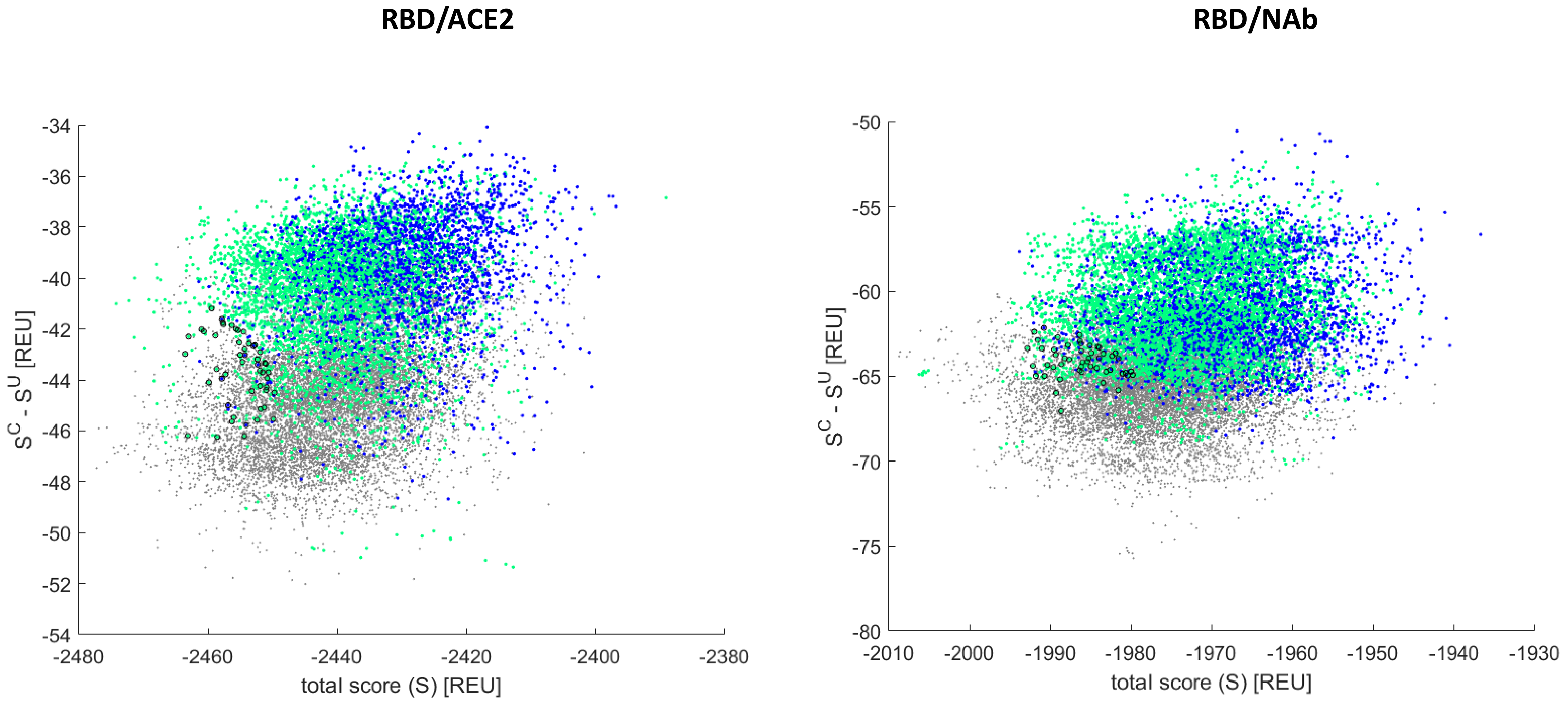
RBD/ACE2
RBD/NAb
-34
-36
-38
-40
-42
-44
-46
-48
-50
-52
-54
S^C - S^U [REU]
-2480
-2460
-2440
-2420
-2400
-2380
total score (S) [REU]
-50
-55
-60
-65
-70
-75
-80
-2010
-2000
-1990
-1980
-1970
-1960
-1950
-1940
-1930

**S13**

### Omicron

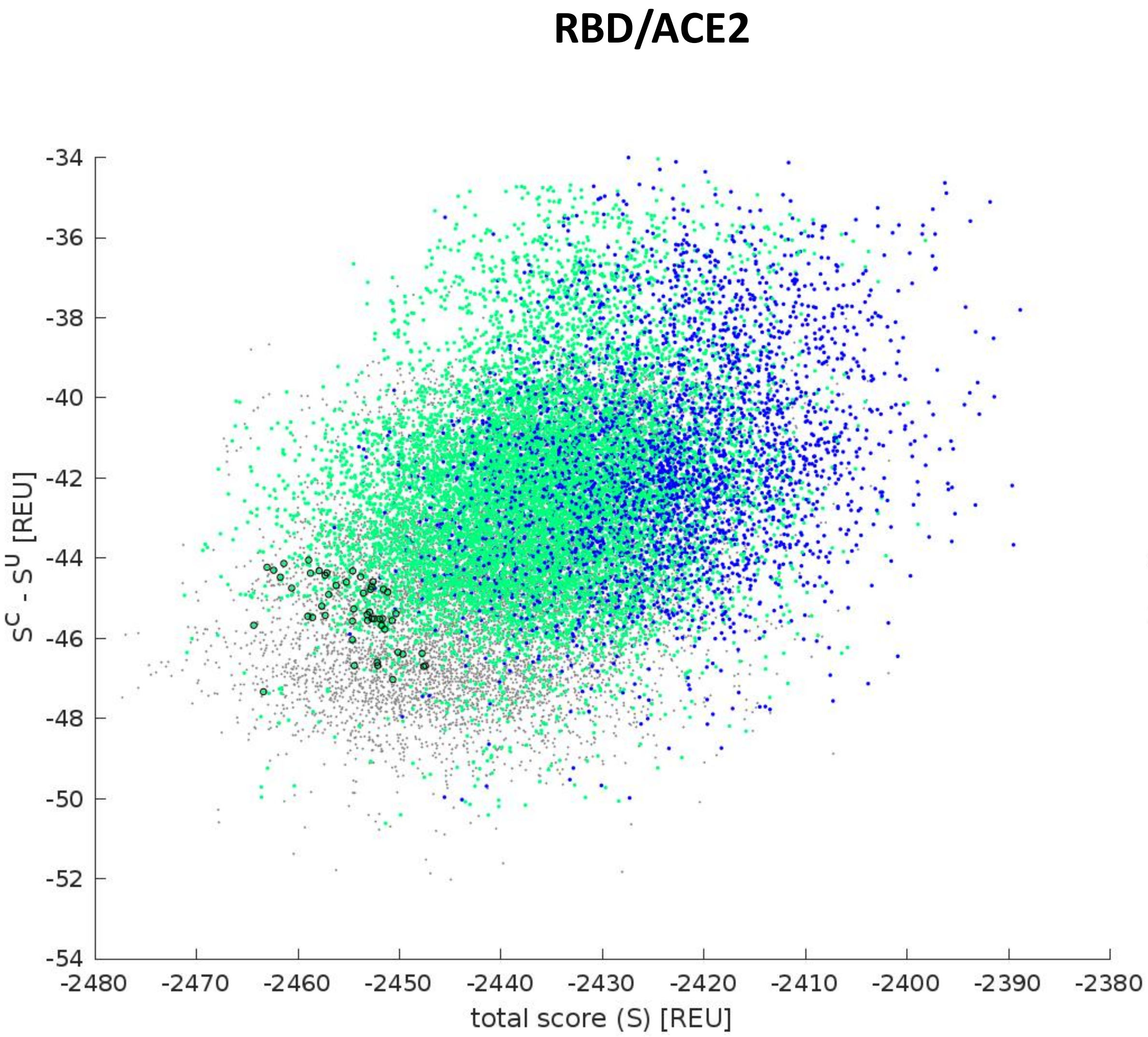
RBD/ACE2
-34
-36
-38
-40
-42
-44
-46
-48
-50
-52
-54
$S^C - S^U$ [REU]
-2480
-2470
-2460
-2450
-2440
-2430
-2420
-2410
-2400
-2390
-2380
total score (S) [REU]

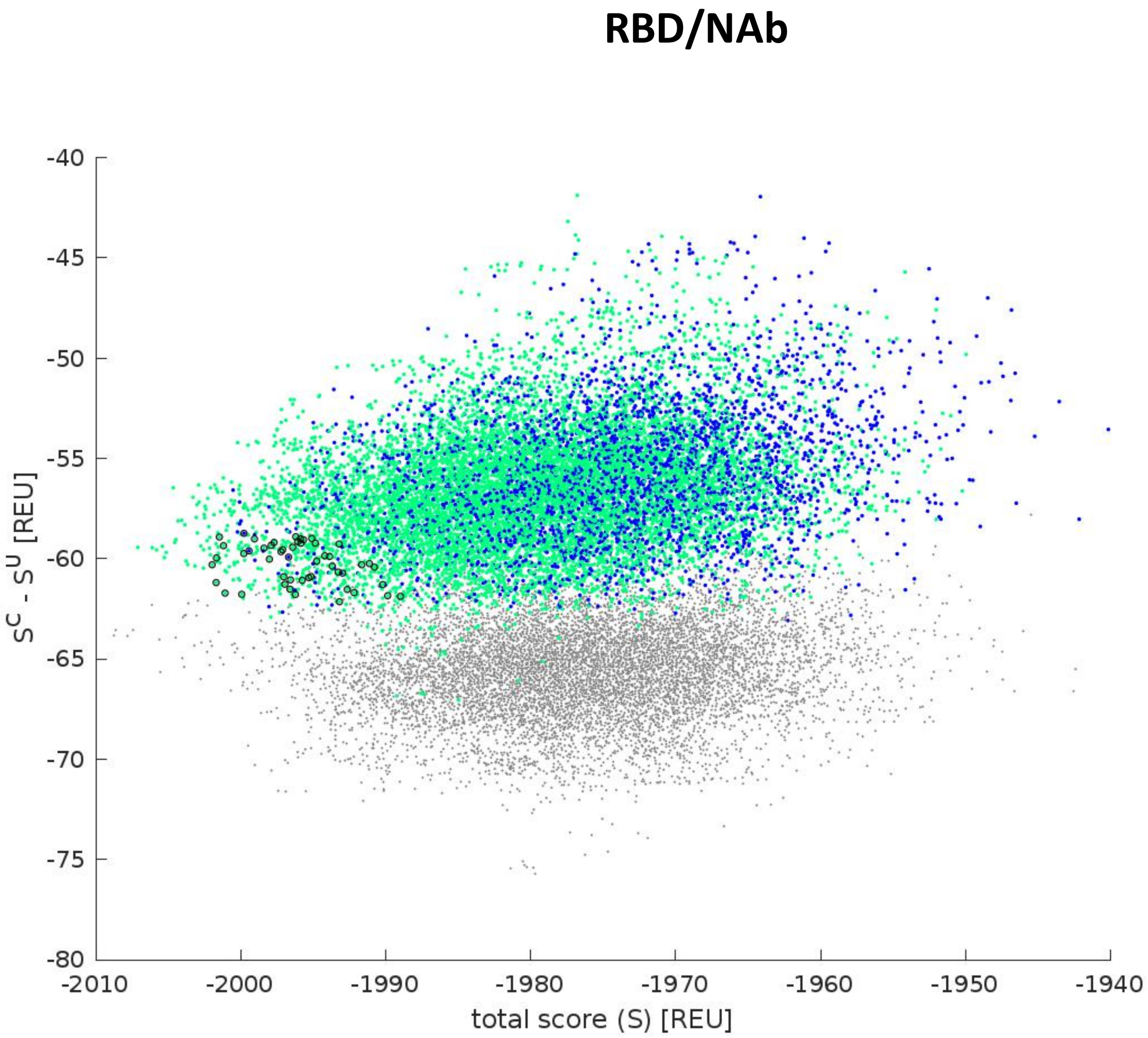
RBD/NAb
-40
-45
-50
-55
-60
-65
-70
-75
-80
$S^C - S^U$ [REU]
-2010
-2000
-1990
-1980
-1970
-1960
-1950
-1940
total score (S) [REU]

**S14**

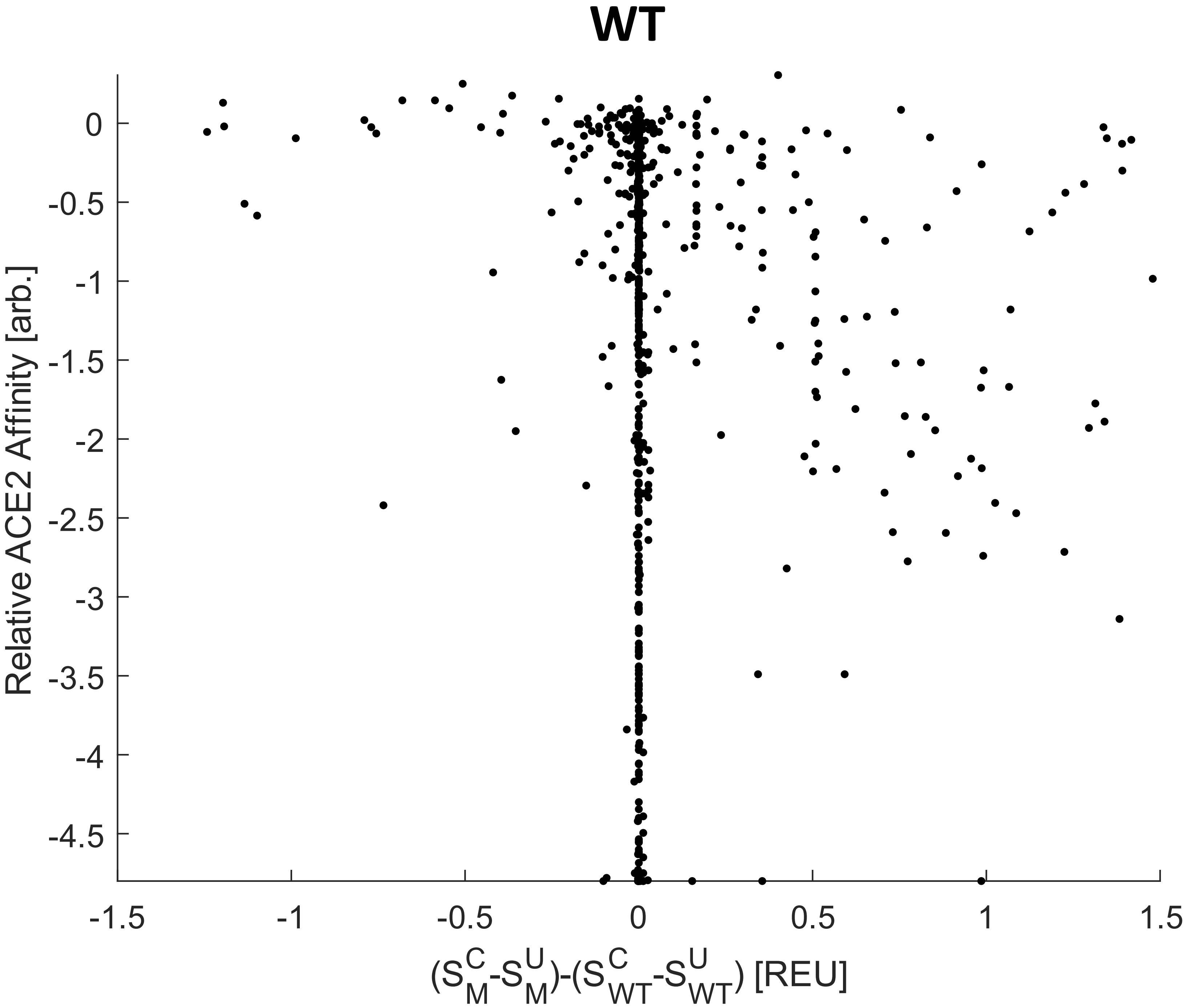
WT
Relative ACE2 Affinity [arb.]
0
-0.5
-1
-1.5
-2
-2.5
-3
-3.5
-4
-4.5
-1.5
-1
-0.5
0
0.5
1
1.5
$(S_M^C - S_M^U) - (S_{WT}^C - S_{WT}^U)$ [REU]

**S15**

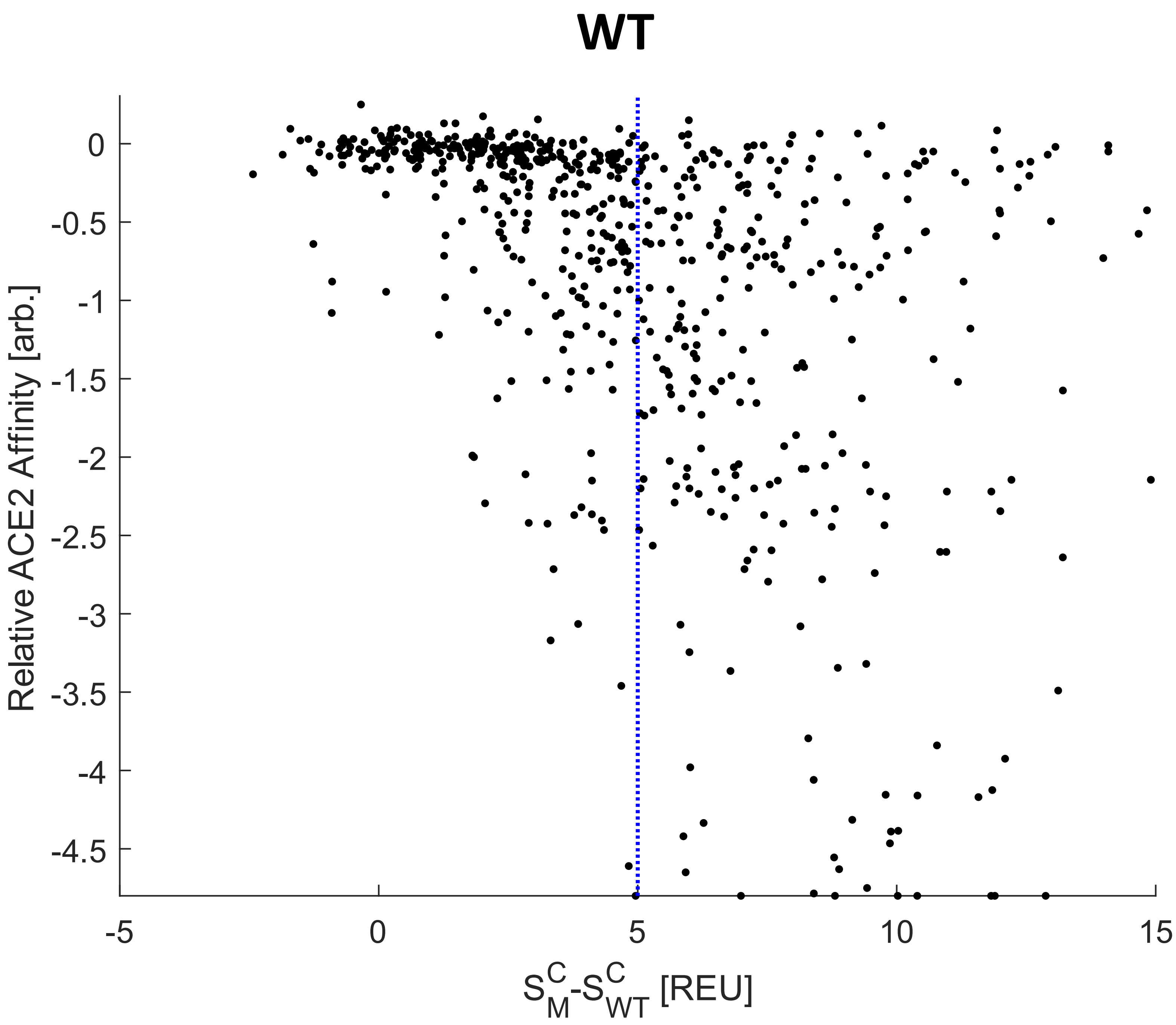
WT
Relative ACE2 Affinity [arb.]
0
-0.5
-1
-1.5
-2
-2.5
-3
-3.5
-4
-4.5
-5
0
5
10
15
$S^C_M$-$S^C_{WT}$ [REU]

**S16**

### Delta

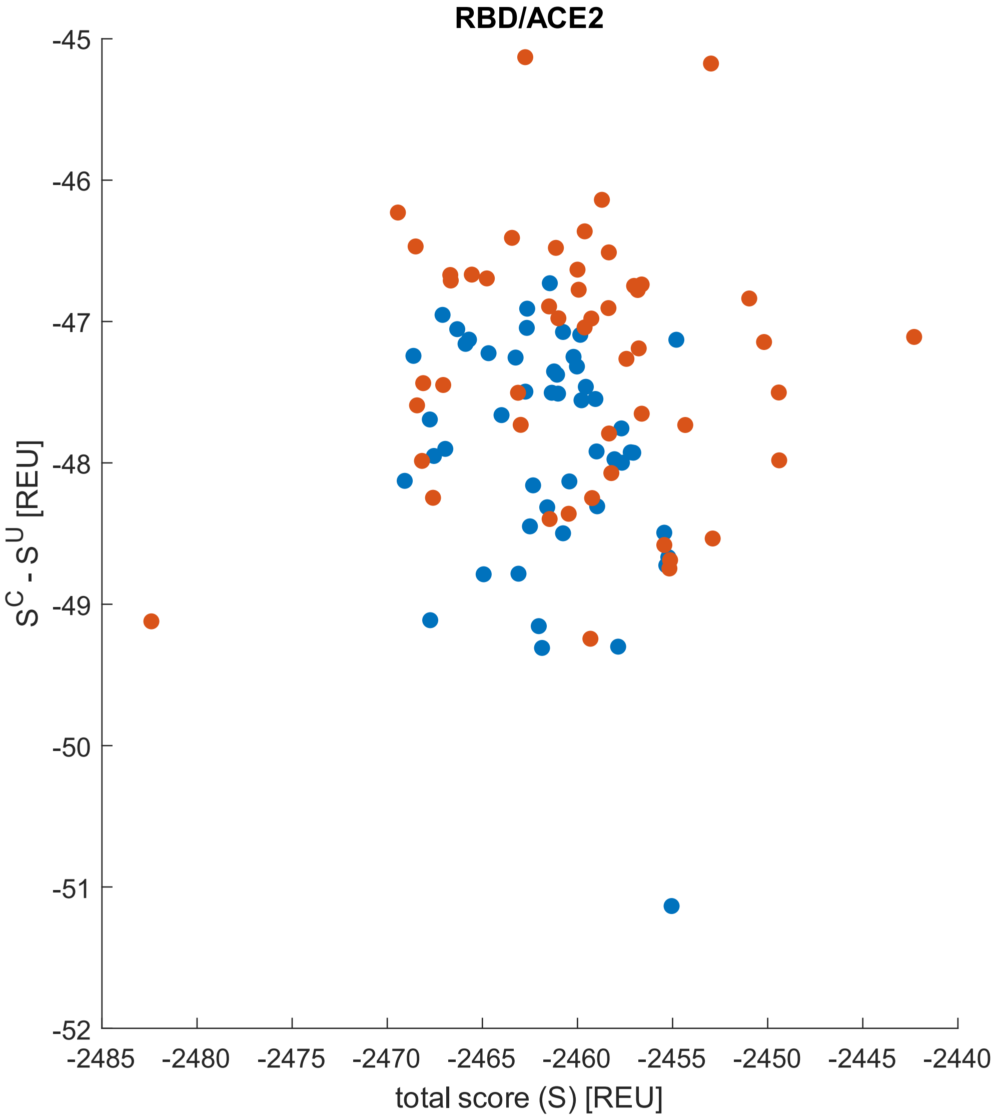

RBD/ACE2
-45
-46
-47
-48
-49
-50
-51
-52
$S^C - S^U$ [REU]
-2485
-2480
-2475
-2470
-2465
-2460
-2455
-2450
-2445
-2440
total score (S) [REU]


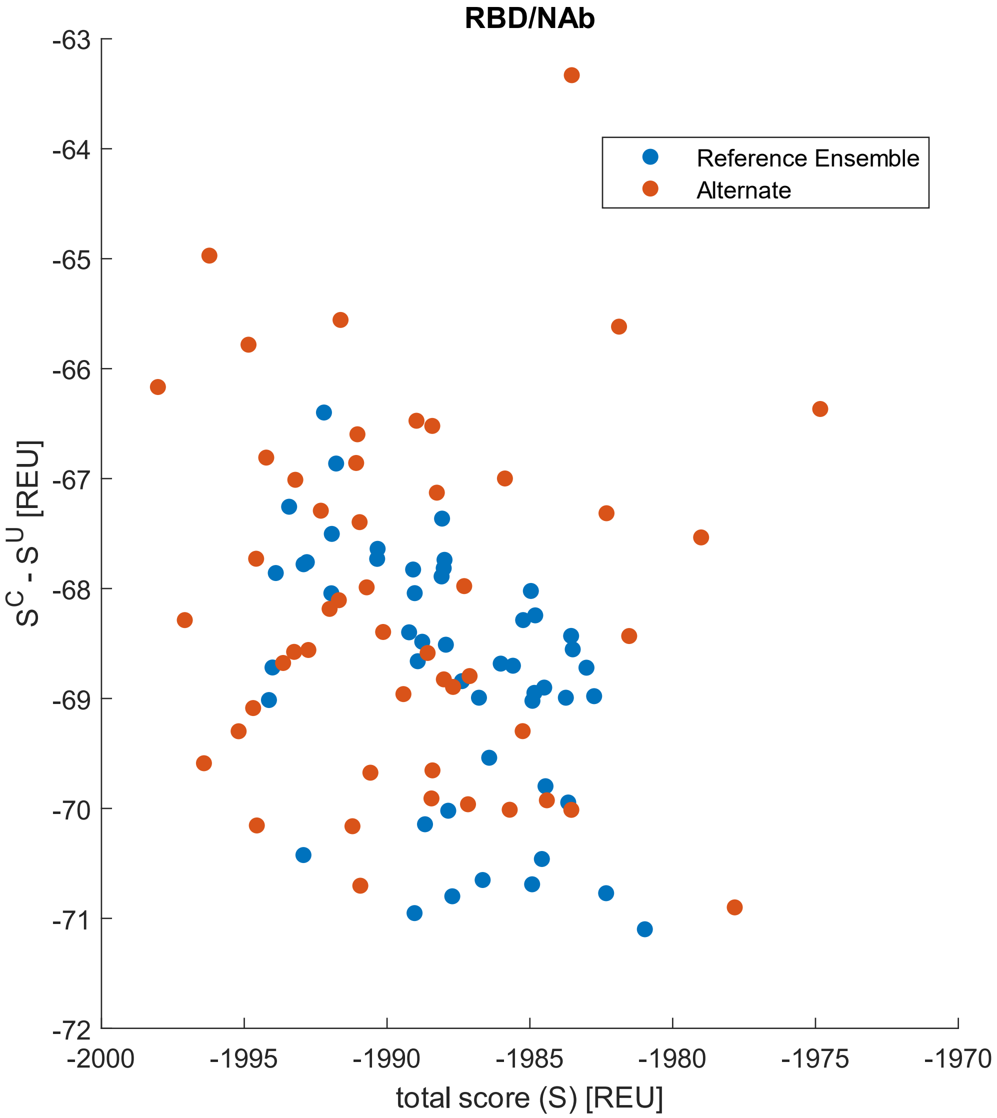

RBD/NAb
Reference Ensemble
Alternate
-63
-64
-65
-66
-67
-68
-69
-70
-71
-72
$S^C - S^U$ [REU]
-2000
-1995
-1990
-1985
-1980
-1975
-1970
total score (S) [REU]

S17

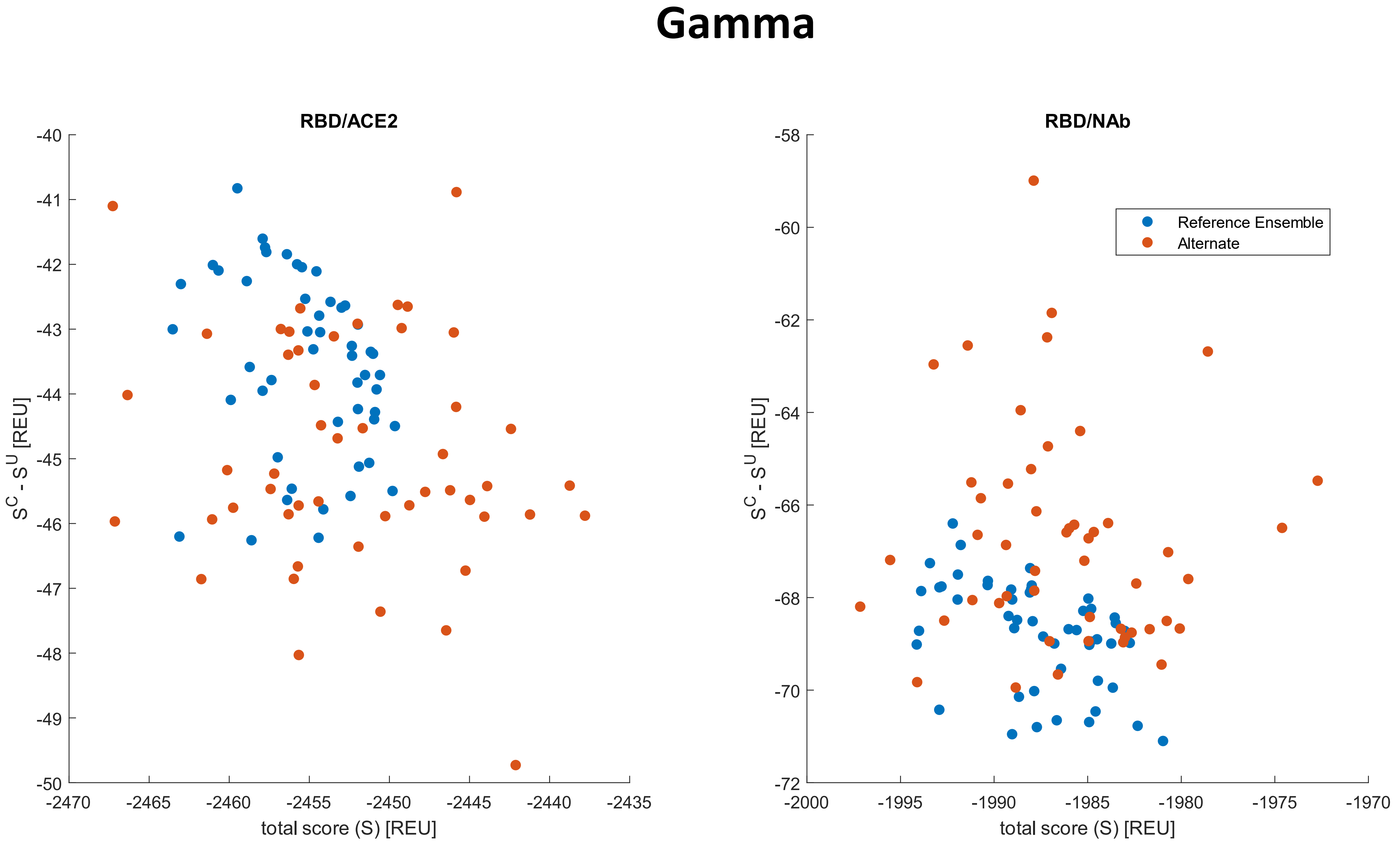
Gamma
RBD/ACE2
$S^C - S^U$ [REU]
total score (S) [REU]
-40
-41
-42
-43
-44
-45
-46
-47
-48
-49
-50
-2470
-2465
-2460
-2455
-2450
-2445
-2440
-2435
RBD/NAb
-58
-60
-62
-64
-66
-68
-70
-72
-2000
-1995
-1990
-1985
-1980
-1975
-1970
Reference Ensemble
Alternate

S18

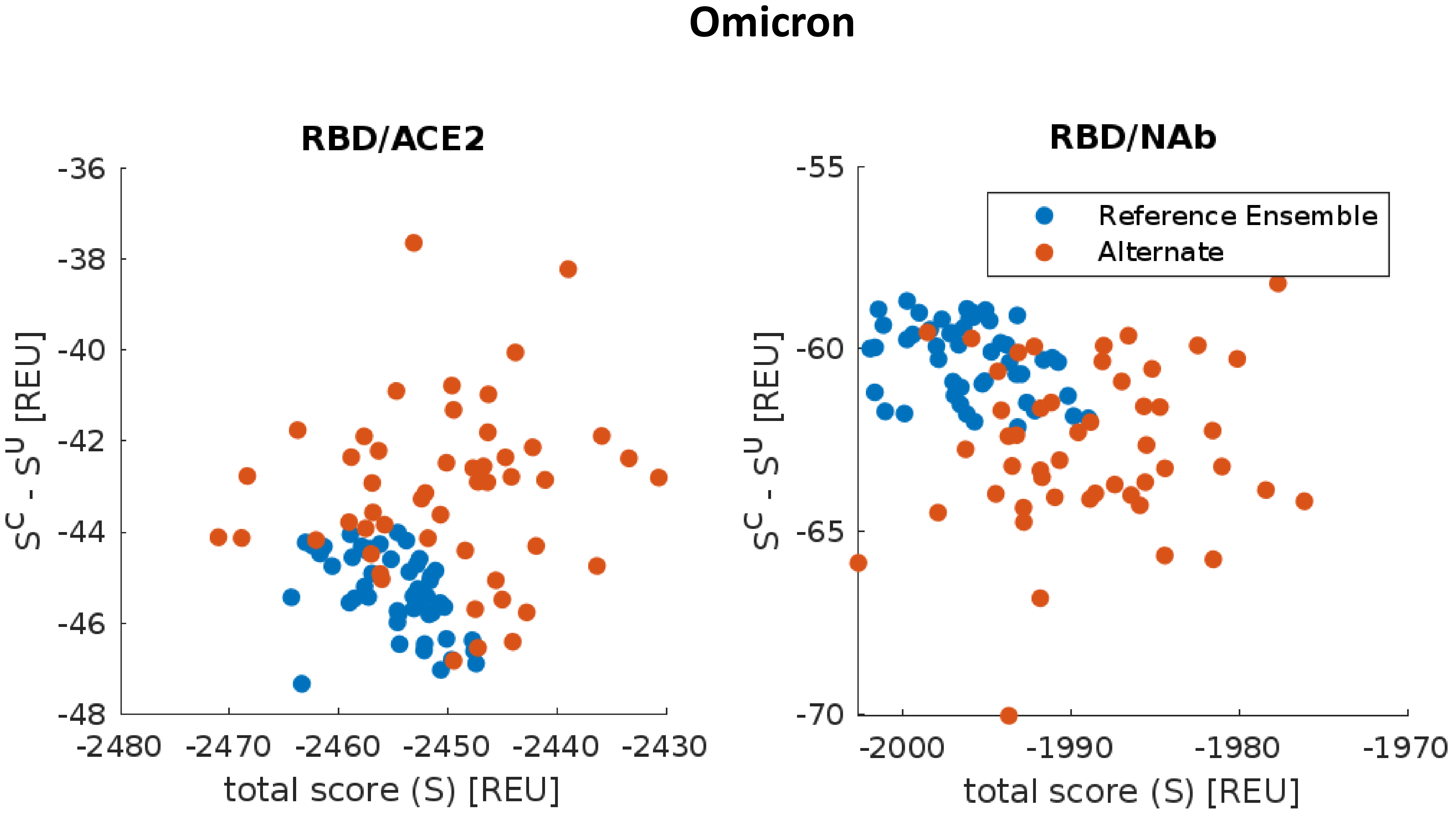
Omicron
RBD/ACE2
-36
-38
-40
-42
-44
-46
-48
-2480
-2470
-2460
-2450
-2440
-2430
$S^c - S^u$ [REU]
total score (S) [REU]
RBD/NAb
-55
-60
-65
-70
-2000
-1990
-1980
-1970
Reference Ensemble
Alternate
$S^c - S^u$ [REU]
total score (S) [REU]

### Table S1

1) fastRelaxSpecRep.xml, which includes backbone minimization and sidechain repacking, is conducted using the following .xml file. This is applied to the WT crystal structure; to the crystal structure after the variant mutations have been introduced; and to the WT conformational ensemble after the variant mutations have been introduced to generate the alternate variant conformations. 5 repetitions are conducted in the below example command line.

Example Command Line:

.../rosetta_scripts.static.linuxgccrelease -parser:protocol .../fastRelaxSpecRep.xml -parser:script_vars numRep=5 -out:suffix suffix -out:path:pdb .../outFolder -out:path:score /dev/null -s .../input.pdb

---

```
<ROSETTASCRIPTS>
	<SCOREFXNS>
	</SCOREFXNS>
	<FILTERS>
	</FILTERS>
	<TASKOPERATIONS>
	</TASKOPERATIONS>
	<MOVERS>
		This mover does a fast relaxation. The input complex is should be pre-relaxed (prior to mutation).
		<FastRelax name="relax" disable_design="1" repeats="%%numRep%%"/>
	</MOVERS>
	<PROTOCOLS>
		Fast-relax the mutated complex.
		<Add mover_name="relax"/>
	</PROTOCOLS>
```

</ROSETTASCRIPTS>

---

2) Example resfile which is used to introduce mutations without repacking other residues. This example modifies site 403 on chain E to be Alanine(A).

---

NATRO

USE_INPUT_SC

start

403 E PIKAA A EX 1 EX 2 EX 3 EX 4

---

3) mutAndPack.xml, which introduces a mutation specified by the resfile. Note that (as in the above example)

if the resfile does not specify sidechain repacking, only the mutation is introduced with the remaining residues unchanged.

Example Command Line:

.../rosetta_scripts.static.linuxgccrelease -parser:protocol .../mutAndPack.xml -parser:script_vars resfile_path=.../resfile.dat -out:suffix suffix -out:path:pdb .../outFolder -out:path:score /dev/null -s input.pdb

---

<ROSETTASCRIPTS>

<SCOREFXNS>

</SCOREFXNS>

<FILTERS>

</FILTERS>

<TASKOPERATIONS>

This task operation is used to specify the location of the resfile to use for design. Note this can also be used to specify "dummy design" X->X

<ReadResfile name="resfile" filename="%%resfile_path%%"/>

</TASKOPERATIONS>

<MOVERS>

This mover packs rotamers with design as specified in the resfile named generic.resfile per the above.

<PackRotamersMover name="mut_and_pack" task_operations="resfile"/>

</MOVERS>

<PROTOCOLS>

First mutate the complex

<Add mover_name="mut_and_pack"/>

</PROTOCOLS>

</ROSETTASCRIPTS>

---

4) runInterfaceDDGandRMSD.xml which may be used to calculate the interface energy; identify interface residues;

and calculate the interface RMSD relative to a reference structure over a comma separated list of residues.

Example Command Line:

.../rosetta_scripts.static.linuxgccrelease -parser:protocol .../runInterfaceDDGandRMSD.xml -parser:script_vars pSep=1 chain1=A chain2=E interString=comma_separated_list_of_interface_residues pIn=0 interface_ID=A_E pUnbound=1 pBound=0 rUnbound=0 rBound=0 chainstomove=E -in:file:native .../reference.pdb -out:file:score_only .../score.txt -s input.pdb

---

```
<ROSETTASCRIPTS>
	<SCOREFXNS>
		<ScoreFunction name="standardfxn" weights="ref2015"/>
	</SCOREFXNS>
	<RESIDUE_SELECTORS>
		<Chain name="chA" chains="%%chain1%%" />
		<Chain name="chB" chains="%%chain2%%" />
		InterfaceByVector name="interfaceAB" grp1_selector="chA" grp2_selector="chB"
		<Index name="interfaceAB" resnums="%%interString%%"
error_on_out_of_bounds_index="1" reverse="0" />
	</RESIDUE_SELECTORS>
	<MOVE_MAP_FACTORIES>
		<MoveMapFactory name="Interface">
	<Backbone residue_selector="interfaceAB"/>
	<Chi residue_selector="interfaceAB"/>
		</MoveMapFactory>
	</MOVE_MAP_FACTORIES>
	<SIMPLE_METRICS>
		 SelectedResidueCountMetric name="nres_int" residue_selector="interfaceAB"
		<RMSDMetric name="Irmsd" rmsd_type="rmsd_protein_bb_ca"
residue_selector="interfaceAB" use_native="1" super="1"/>
		<RMSDMetric name="chB_rmsd" rmsd_type="rmsd_protein_bb_ca"
residue_selector="chB" use_native="1" super="1"/>
		<RMSDMetric name="chA_rmsd" rmsd_type="rmsd_protein_bb_ca"
residue_selector="chA" use_native="1" super="1"/>
	</SIMPLE_METRICS>
	<MOVERS>
		This mover applies the total score
		<ScoreMover name="apply_score" scorefxn="standardfxn"/>
```

This mover calculates the interface energy.

<InterfaceAnalyzerMover name="calc_interface" pack_separated="%%pSep%%" pack_input="%%pIn%%" interface_sc="1" interface="%%interface_ID%%"/>

This mover does a fast relaxation. The input complex is should be pre-relaxed (prior to mutation).

<FastRelax name="relax" disable_design="1"/>

This mover calculates DDG

<ddG name="standard_ddG_mover" scorefxn="standardfxn" relax_mover="relax" repack_unbound="%%pUnbound%%" repack_bound="%%pBound%%" relax_bound="%%rBound%%" relax_unbound="%%rUnbound%%" translate_by="1000.0" chain_name="%%chainstomove%%"/>

<RunSimpleMetrics name="run_metrics1" metrics="Irmsd" prefix="I_"/>

<RunSimpleMetrics name="run_metrics2" metrics="chB_rmsd" prefix="chB_"/>

<RunSimpleMetrics name="run_metrics3" metrics="chA_rmsd" prefix="chA_"/>

</MOVERS>

<PROTOCOLS>

First get total score

<Add mover_name="apply_score"/>

Then do interface calculation

<Add mover_name="calc_interface"/>

Then calc ddg

<Add mover_name="standard_ddG_mover"/>

Finally calc RMSD

<Add mover_name="run_metrics1"/>

<Add mover_name="run_metrics2"/>

<Add mover_name="run_metrics3"/>

</PROTOCOLS>

</ROSETTASCRIPTS>

---

5)runInterfaceAndDDG.xml, alternative to above when RMSD calculation is not necessary.

Example Command Lines:

Returns Only Scores:

.../rosetta_scripts.static.linuxgccrelease -parser:protocol .../runInterfaceAndDDG.xml -parser:script_vars pSep=0 pIn=0 interface_ID=A_E pUnbound=0 pBound=0 rUnbound=0 rBound=0 chainstomove=E -out:file:score_only output.txt -s input.pdb

Returns Only PDBs with Interface Residues Specified:

.../rosetta_scripts.static.linuxgccrelease -parser:protocol ../rosetta/my_xml/runInterfaceAndDDG.xml -parser:script_vars pSep=0 pIn=0 interface_ID=HL_E pUnbound=0 pBound=0 rUnbound=0 rBound=0 chainstomove=E -out:path:pdb .../outFolder -out:path:score /dev/null -s input.pdb

---

```
<ROSETTASCRIPTS>
	<SCOREFXNS>
		<ScoreFunction name="standardfxn" weights="ref2015"/>
	</SCOREFXNS>
	<FILTERS>
	</FILTERS>
	<TASKOPERATIONS>
	</TASKOPERATIONS>
	<MOVERS>
		This mover applies the total score
		<ScoreMover name="apply_score" scorefxn="standardfxn"/>
```

This mover calculates the interface energy.

<InterfaceAnalyzerMover name="calc_interface" pack_separated="%%pSep%%" pack_input="%%pIn%%" interface_sc="1" interface="%%interface_ID%%"/>

This mover does a fast relaxation. The input complex is should be pre-relaxed (prior to mutation).

<FastRelax name="relax" disable_design="1"/>

This mover calculates DDG

<ddG name="standard_ddG_mover" scorefxn="standardfxn" relax_mover="relax" repack_unbound="%%pUnbound%%" repack_bound="%%pBound%%" relax_bound="%%rBound%%" relax_unbound="%%rUnbound%%" translate_by="1000.0" chain_name="%%chainstomove%%"/>

</MOVERS>

<PROTOCOLS>

First get total score

<Add mover_name="apply_score"/>

Then do interface calculation

<Add mover_name="calc_interface"/>

Then calc ddg

<Add mover_name="standard_ddG_mover"/>

</PROTOCOLS>

</ROSETTASCRIPTS>
